# scROMA: batch-aware pathway-activity inference and a ground-truth simulation framework for single-cell transcriptomics

**DOI:** 10.64898/2026.08.07.743516

**Authors:** Altynbek Zhubanchaliyev, Matthieu Najm, Victor Laigle, Eric Bonnet, Loredana Martignetti

## Abstract

**Background:** Pathway-activity analysis summarizes gene-level single-cell measurements into interpretable functional modules, but widely used methods lack an integrated significance framework, do not account for the batch effects that pervade multi-sample studies, and are not natively interoperable with Python-based workflows. The field also lacks simulation resources with ground-truth pathway activity for quantitative benchmarking.

**Results:** We present scROMA, a singular-value-decomposition-based method that quantifies pathway activity as coordinated variation, with per-cell scores, per-gene contributions, and permutation-based significance, natively integrated with the Scanpy/AnnData ecosystem. Its batch-aware extension is, to our knowledge, the first to correct batch effects within the gene-set subspace rather than across the full transcriptome, isolating technical variation at the pathway level while preserving signal in other genes. We also release a generative simulation framework producing synthetic data with fully specified ground-truth activities. On simulated benchmarks scROMA is competitive across tasks, and under batch effects its batch-aware mode recovers ordinal pathway structure that full-transcriptome integration misses. Across cystic fibrosis airway, intestinal-organoid, breast cancer, and lung cancer datasets it recovers established biology while separating it from technical and inter-donor variation; in the intestinal-organoid atlas it reproducibly recovers an inflammatory program across donors, separates its sustained from transient components, and resolves cell-type-specific niche-factor targets.

**Conclusions:** scROMA is open-source and released with the simulation framework and pre-generated benchmark datasets as a community resource, providing a scalable, statistically grounded, and batch-aware approach to pathway-level analysis in single-cell transcriptomics.

## 1 Background

Single-cell RNA sequencing (scRNA-seq) has transformed our ability to dissect cellular heterogeneity across tissues, developmental stages, and disease states [1–3]. The rapid growth of single-cell atlases, now encompassing millions of cells from diverse organs, species, and pathological conditions [4–6], has shifted the analytical bottleneck from data generation to biological interpretation. While differential expression analysis identifies individual genes that distinguish cell populations, it provides limited insight into the coordinated transcriptional programs that define cellular function. Pathway activity analysis addresses this limitation by aggregating gene-level signals into biologically coherent units, enabling researchers to characterize cell types and states in terms of the processes they engage rather than the individual genes they express [7, 8].

Several computational approaches have been developed to score pathway activity in single-cell data, each with distinct strengths and limitations. A systematic benchmark by Zhang et al. [9] identified Pagoda2 [10] as achieving the strongest overall performance among early methods; Pagoda2 leverages the first principal component (PC1) of each gene set to quantify pathway overdispersion, but produces a single pathway-level statistic rather than per-cell activity scores, precluding its use for downstream analyses that require single-cell resolution. AUCell [11], one of the most widely adopted methods, scores gene set activity by evaluating the enrichment of pathway genes among the top-ranked genes within each cell. While AUCell operates at single-cell resolution, it does not provide a statistical framework for distinguishing biologically active pathways from inactive ones, a distinction that is critical when screening large gene set collections. Gene Set Enrichment Analysis (GSEA) [7], adapted for single-cell use through frameworks such as Decoupler [12], can identify significantly enriched pathways, but its ranking-based approach was not designed for the sparse, zero-inflated expression profiles characteristic of scRNA-seq data. Scanpy’s `score_genes` [13], the Python equivalent of Seurat’s AddModuleScore [14], provides a computationally efficient per-cell score but similarly lacks a built-in significance framework. More recent methods, including GSDensity [15], SCPA [16], and scGSEA [17], have introduced pathway-centric analyses tailored to single-cell data. Although several of these methods provide per-cell scores or statistical testing individually, SVD-based approaches, which captured coordinated gene set variability most effectively in prior benchmarks [9], have not yet been extended to combine single-cell resolution activity scores with a permutation-based significance framework within the Scanpy/AnnData ecosystem [13] that supports a large proportion of current single-cell workflows.

A second, largely unaddressed challenge concerns the handling of batch effects in pathway activity inference. Modern single-cell studies routinely integrate data across patients, institutions, and sequencing platforms, introducing pervasive technical variation that can confound pathway-level signals [18, 19]. In current practice, batch correction methods such as Harmony [20], scVI [21], or ComBat [22] are most commonly applied to define shared cell populations across batches, after which downstream analyses such as differential expression testing and pathway enrichment are performed on the uncorrected expression matrix, often including batch as a covariate in statistical models. Some studies, however, compute pathway scores directly from the corrected expression matrix [23]. This latter approach carries a structural limitation: global batch correction removes technical variance across all genes simultaneously and can inadvertently attenuate biological signal, particularly for gene sets whose variation is correlated with, but distinct from, the batch axis. Indeed, recent work has shown that full-transcriptome batch correction can introduce spurious expression patterns even in the absence of true batch effects [24]. Importantly, neither the covariate-based nor the corrected-matrix approach operates at the level of the gene set itself, meaning that technical variance affecting a specific pathway’s constituent genes cannot be isolated and removed without altering the broader expression landscape. This is consequential because batch effects vary in magnitude across genes and can disproportionately affect specific functional categories [25, 26]. To our knowledge, no existing pathway scoring method integrates batch correction directly within the gene-set subspace.

A third limitation addressed in this work is the absence of rigorous benchmarking infrastructure. Evaluating pathway activity methods requires datasets with known ground-truth pathway activation states at single-cell resolution, information that real datasets, by definition, cannot provide. Existing single-cell simulation frameworks such as Splatter [27] and scDesign3 [28] are designed for benchmarking differential expression, clustering, or batch correction methods, and do not model pathway-level activation structures. As a result, the field has relied primarily on indirect validation through biological plausibility, limiting the ability to make quantitative, head-to-head comparisons between methods.

To address these challenges, we developed scROMA, a scalable Python implementation of the ROMA (Representation and Quantification of Module Activity) algorithm [29] for single-cell transcriptomics, natively integrated with the Scanpy/AnnData ecosystem. scROMA infers pathway activity from coordinated variation within gene-set expression submatrices, assesses statistical significance, and introduces a batch-aware extension that corrects technical variation directly within the gene-set subspace. To enable quantitative benchmarking, we additionally developed a generative simulation framework that produces synthetic scRNA-seq count matrices with fully specified ground-truth pathway activities at single-cell resolution, together with three complementary evaluation tasks that assess methods at progressively finer granularity.

We validated scROMA through a series of analyses providing converging evidence across controlled and real-world settings: (i) fully controlled simulations with known ground truth; (ii) a cystic fibrosis airway epithelium atlas [31]; (iii) a human intestinal-organoid niche-factor perturbation atlas [40]; (iv) a breast cancer tumor microenvironment atlas [32]; and (v) a lung cancer dataset annotated with inferred copy-number profiles [33]. scROMA is open-source software and is released together with the simulation framework and pre-generated benchmark datasets as a community resource.

## 2 Results

### 2.1 scROMA: SVD-based pathway activity scoring with batch-aware extension

We developed scROMA, a Python implementation of the ROMA (Representation and Quantification Of Module Activity) algorithm [29] for quantifying pathway activity in single-cell transcriptomics data, natively integrated with the Scanpy/AnnData ecosystem [13] (Fig. 1). For each gene set, scROMA subsets the expression matrix to the member genes and performs singular value decomposition (SVD) on the transposed submatrix (genes × cells). The decomposition yields two complementary pathway-level summary statistics: the L1 score, which quantifies the degree to which expression variation within the gene set is concentrated along a single axis (overdispersion), and the median expression score, which captures systematic up- or down-regulation of the pathway (shift). These two modes are complementary: overdispersion captures coordinated variability, whereas shift captures coherent directional change. In addition, the decomposition provides per-cell activity scores (projections onto the first principal component) at single-cell resolution and fully interpretable gene-level contributions identifying which genes drive the pathway activity signal. Outlier genes whose removal disproportionately affects the L1 score are detected and excluded via leave-one-out cross-validation, and statistical significance is assessed against a permutation-derived null distribution with Benjamini-Hochberg correction [36] for multiple testing (see Methods for full algorithmic details).

**Fig. 1:**
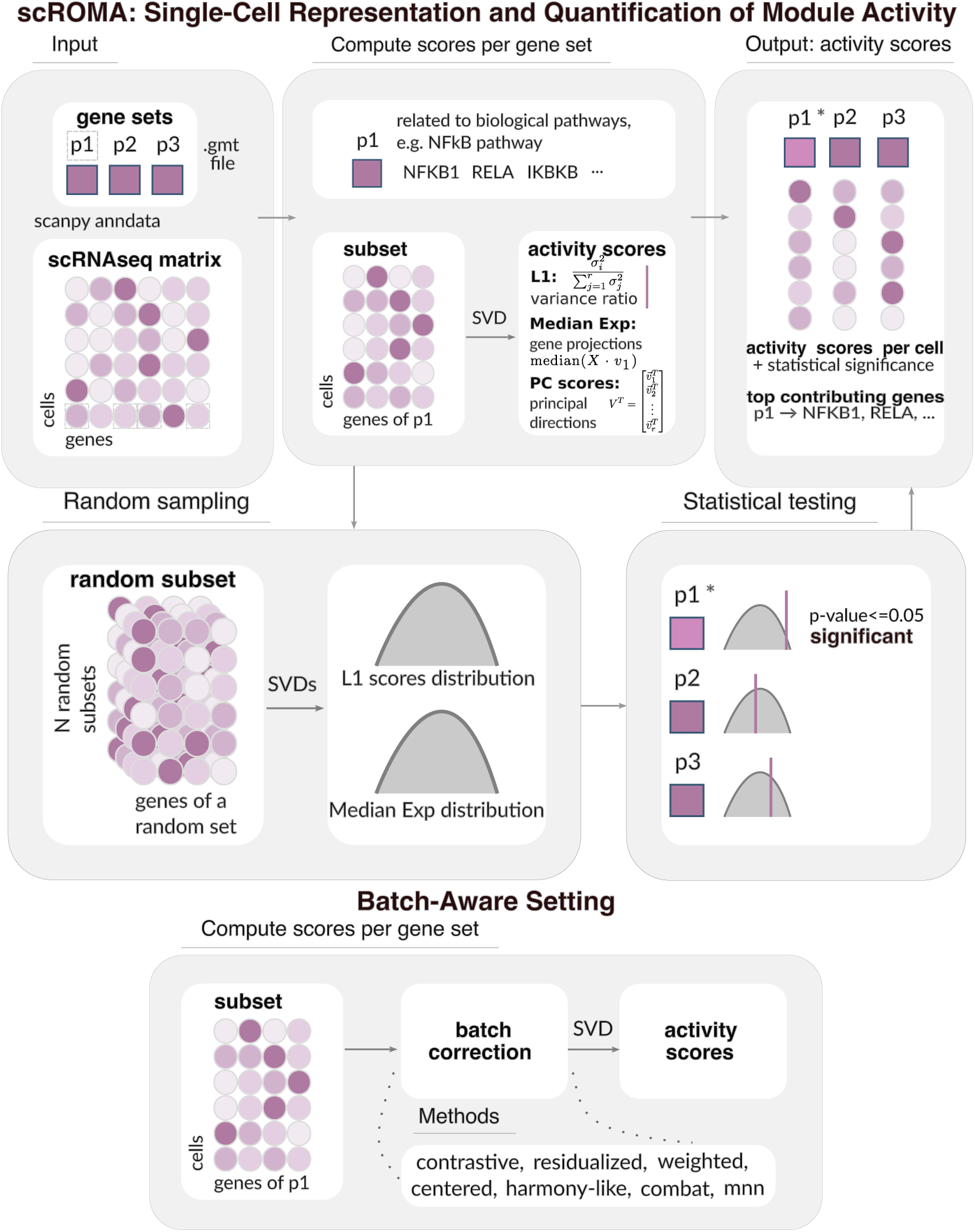
Overview of the scROMA algorithm. For each gene set, scROMA subsets the expression matrix to member genes and performs SVD on the transposed submatrix (genes × cells), yielding two complementary pathway-level statistics (L1 overdispersion score and median expression score), per-cell activity scores, and gene-level contribution weights. Statistical significance is assessed via a permutation-based null distribution. The batch-aware extension (bottom) integrates batch correction directly within the gene-set subspace prior to SVD, rather than correcting the full expression matrix.

To enable analysis of multi-batch datasets, we introduce a batch-aware extension that is, to our knowledge, the first pathway scoring method to integrate batch correction directly within the gene-set subspace (Fig. 1, bottom). Rather than applying batch correction to the full expression matrix prior to pathway scoring — the conventional approach, which can inadvertently attenuate biological signal — the batch-aware extension applies correction within the gene-set-level submatrix before SVD, removing technical confounding at the pathway level while preserving biological variation encoded in genes outside the queried gene set. Seven correction strategies are available, spanning a spectrum from linear adjustments (residualized, contrastive, weighted, and centered PCA) to more aggressive methods (ComBat [22], Harmony-like iterative centroid alignment [20], and mutual nearest neighbors [34]), providing flexibility for different batch effect structures (see Methods). The scROMA software natively accepts AnnData objects and stores all results within the AnnData structure for seamless integration with downstream Scanpy workflows.

### 2.2 A simulation framework for evaluating pathway activity inference methods

To enable rigorous, quantitative benchmarking of pathway activity inference methods, we developed a generative simulation framework that produces synthetic scRNA-seq count matrices with fully known ground-truth pathway activation states at single-cell resolution (Fig. 2A). The framework captures key features of real scRNA-seq data — including count-based noise, expression-dependent dropout, library size heterogeneity, and batch effects — while providing exact ground truth for pathway activity. We release both the simulation code and pre-generated benchmark datasets as a community resource [46, 49].

**Fig. 2:**
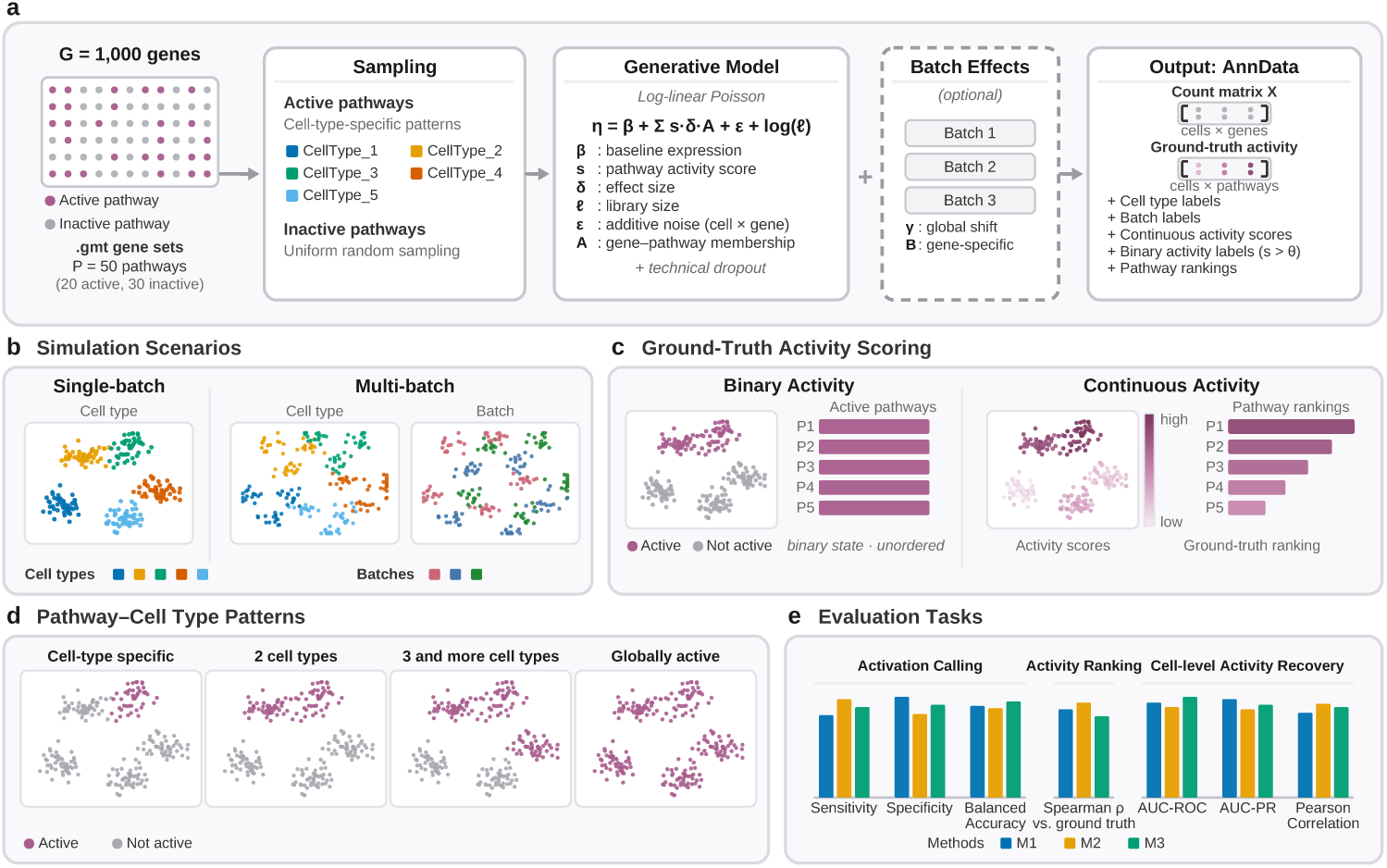
Generative simulation framework for benchmarking pathway activity methods. **(a)** Overview of the simulation pipeline producing synthetic scRNA-seq count matrices with known ground-truth pathway activities. **(b)** Example UMAP embedding of simulated cells colored by cell type and batch for two simulation scenarios. **(c)** Ground-truth pathway activities provided as continuous per-cell scores and binary activation labels. **(d)** Pathway-cell-type activation patterns used as ground truth, spanning four scenarios: cell-type-specific (active in a single cell type), shared between two cell types, shared across three or more cell types, and globally active (present in all cell types). **(e)** Three complementary evaluation tasks: Task 1 (Activation Calling), Task 2 (Activity Ranking), and Task 3 (Cell-Level Activity Recovery).

We defined a gene universe of 1,000 genes and constructed 50 pathway gene sets, of which 20 were designated as active and 30 served as inactive decoys. To capture realistic gene set architectures found in curated databases such as MSigDB [8], we designed three structural blocks among the active pathways with varying degrees of gene sharing: a disjoint block representing cell-type-specific marker programs, a high-overlap block modeling functionally related pathways, and a moderate-overlap block reflecting the heterogeneous overlap typical of curated pathway collections (Fig. 2D; see Methods for the complete specification of pathway architectures, cell-type activation patterns, and all generative model parameters). We simulated five cell types with defined pathway activation patterns spanning four biological scenarios: strictly cell-type-specific pathways, pathways shared between two cell types, globally active pathways present in all cell types, and shared patterns across three or more cell types (Fig. 2D). This combinatorial design provides ground truth for evaluating both per-cell pathway detection and cell-type-level enrichment analyses.

Counts were generated using a log-linear Poisson model incorporating gene-specific baseline expression, pathway activity contributions, additive noise, and library size offsets, with expression-dependent dropout recapitulating the sparsity characteristic of scRNA-seq data (see Methods for model equations and parameter values). Ground-truth pathway activities were provided in two forms: continuous per-cell scores establishing a ground-truth ranking, and binary labels separating genuinely active from background-level cells (Fig. 2C). In the single-batch regime, we simulated 2,000 cells without batch effects; in the multi-batch regime, we simulated three batches with 1,000 cells each, incorporating global batch shifts, gene-specific batch effects, and batch-dependent library sizes while keeping pathway activities identical across batches, ensuring that any degradation in recovery reflects technical confounding rather than biological differences (see Methods).

We designed three complementary evaluation tasks (Fig. 2E). Task 1 — Activation Calling — assesses the ability to correctly distinguish active from inactive pathways, measured by sensitivity, specificity, and balanced accuracy. Task 2 — Activity Ranking — evaluates whether methods preserve the ground-truth ordering of pathway activities, quantified by Spearman *ρ*. Task 3 — Cell-Level Activity Recovery — measures the accuracy of per-cell activity scores via AUC-ROC, AUC-PR, and Pearson correlation. Higher values indicate better performance across all metrics. Simulated datasets are provided as AnnData objects with full ground-truth annotations and all generative parameters [49], and the simulation code supports user-specified custom scenarios — analogous to the role that Splatter [27] plays for scRNA-seq simulation more broadly (see Methods for formal metric definitions and parameter tables).

### 2.3 scROMA is competitive on simulated benchmarks and uniquely robust to batch effects

We benchmarked scROMA against the most widely used single-cell pathway-scoring methods — AUCell [11], GSEA [7], and Scanpy’s `score_genes` [13] — across the three evaluation tasks on 10 replicate simulated datasets with identical generative parameters but different random seeds. In the multi-batch scenario we additionally paired each competitor with scVI [21] integration and evaluated all scROMA batch-aware variants (Fig. 3; see Methods).

**Fig. 3:**
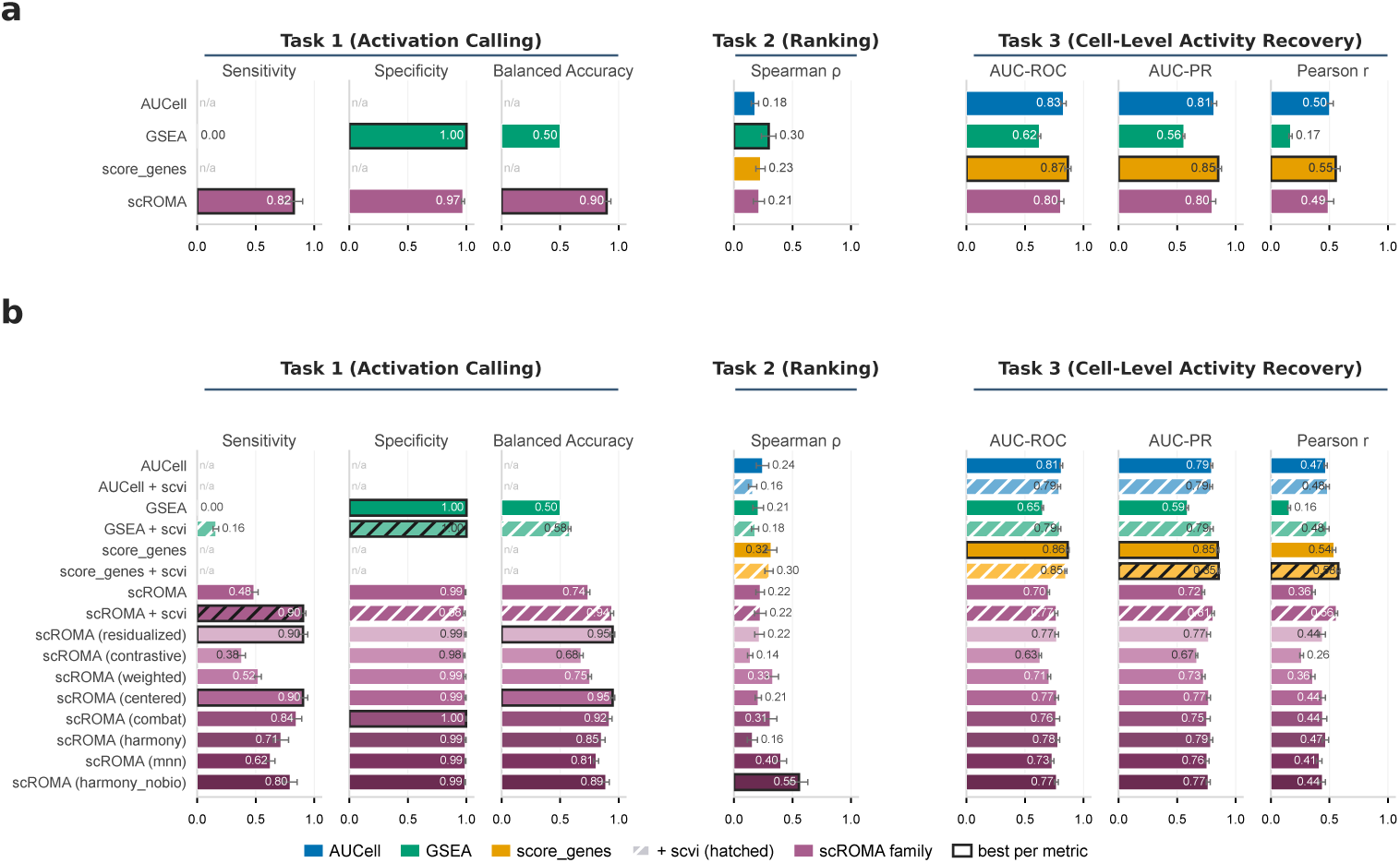
Benchmarking scROMA against existing pathway activity methods on simulated data. Performance comparison across 10 replicate simulated datasets for single-batch **(a)** and multi-batch **(b)** scenarios. Task 1 (Activation Calling): sensitivity, specificity, and balanced accuracy. Task 2 (Activity Ranking): Spearman *ρ* with ground-truth pathway ranking. Task 3 (Cell-Level Activity Recovery): AUC-ROC, AUC-PR, and Pearson correlation with ground-truth per-cell scores. Methods compared: scROMA (base and batch-aware variants), AUCell, GSEA, `score—genes`, and scVI-precorrected alternatives. Bar heights indicate means across replicates; error bars indicate the standard error of the mean.

On activation calling (Task 1), scROMA was the only method that recovered active pathways: without batch effects it reached balanced accuracy 0.90 (sensitivity 0.82, specificity 0.97), whereas GSEA, the only competitor that performs this task, recovered none of them (sensitivity 0.00; balanced accuracy 0.50, rising to 0.58 with scVI), and AUCell and `score_genes` do not support it. Under multi-batch confounding, uncorrected scROMA degraded sharply (balanced accuracy 0.74, sensitivity 0.49), and correcting within the gene-set subspace recovered it: the residualized and centered strategies reached 0.95 (sensitivity 0.91), ComBat 0.92, and the label-free Harmony-like variant 0.89, with the eight batch-aware arms spanning 0.68−0.95. On cell-level activity recovery (Task 3), scROMA performed competitively, with `score_genes` the strongest overall (AUC-ROC 0.86−0.87, AUC-PR 0.85) and scROMA’s best variants close behind (AUC-ROC 0.80 without batch effects and up to 0.78 under them).

The decisive difference emerged in activity ranking under batch effects (Task 2). Without batch effects, all methods recovered ground-truth rankings only modestly (Spearman *ρ* 0.18−0.30). Under multi-batch confounding, however, scROMA’s best batch-aware variant — the label-free Harmony-like correction — reached *ρ* = 0.55 士 0.24 (mean ± s.d. across the ten replicates) against 0.32 ± 0.16 for the strongest competitor (`score_genes`), while pairing competitors with scVI integration did not improve ranking (`score_genes` + scVI, 0.30). scROMA’s pathway-subspace correction thus recovers ordinal structure that full-transcriptome integration misses, without a separate integration step.

A complementary benchmark on a real PBMC IFN-*β* dataset [30] augmented with strong synthetic batch effects — with the interferon response as biological ground truth — gave concordant results, scROMA’s batch-aware mode matching or exceeding the two-step workflow (Supplementary Note 1).

### 2.4 Application to cystic fibrosis airway epithelium identifies subpopulation-specific pathway activities

To evaluate scROMA on a real-world disease atlas, we applied it to a multi-institutional single-cell RNA-seq dataset of proximal airway epithelium from patients with end-stage cystic fibrosis (CF, *n* = 19) and healthy controls (CO, *n* = 19) [31], comprising 40,709 cells across 16 epithelial subpopulations — five basal (Basal1—5), five secretory (Secretory1-5), three ciliated (Ciliated1—3), and three rare types (FOXN4^+^, ionocyte, neuroendocrine) — processed across three institutions with distinct isolation protocols. After scANVI [41] integration to correct institution-level technical variation, scROMA in base mode identified 26 significantly active Hallmark pathways [8] at single-cell resolution (Fig. 4; see Methods).

**Fig. 4:**
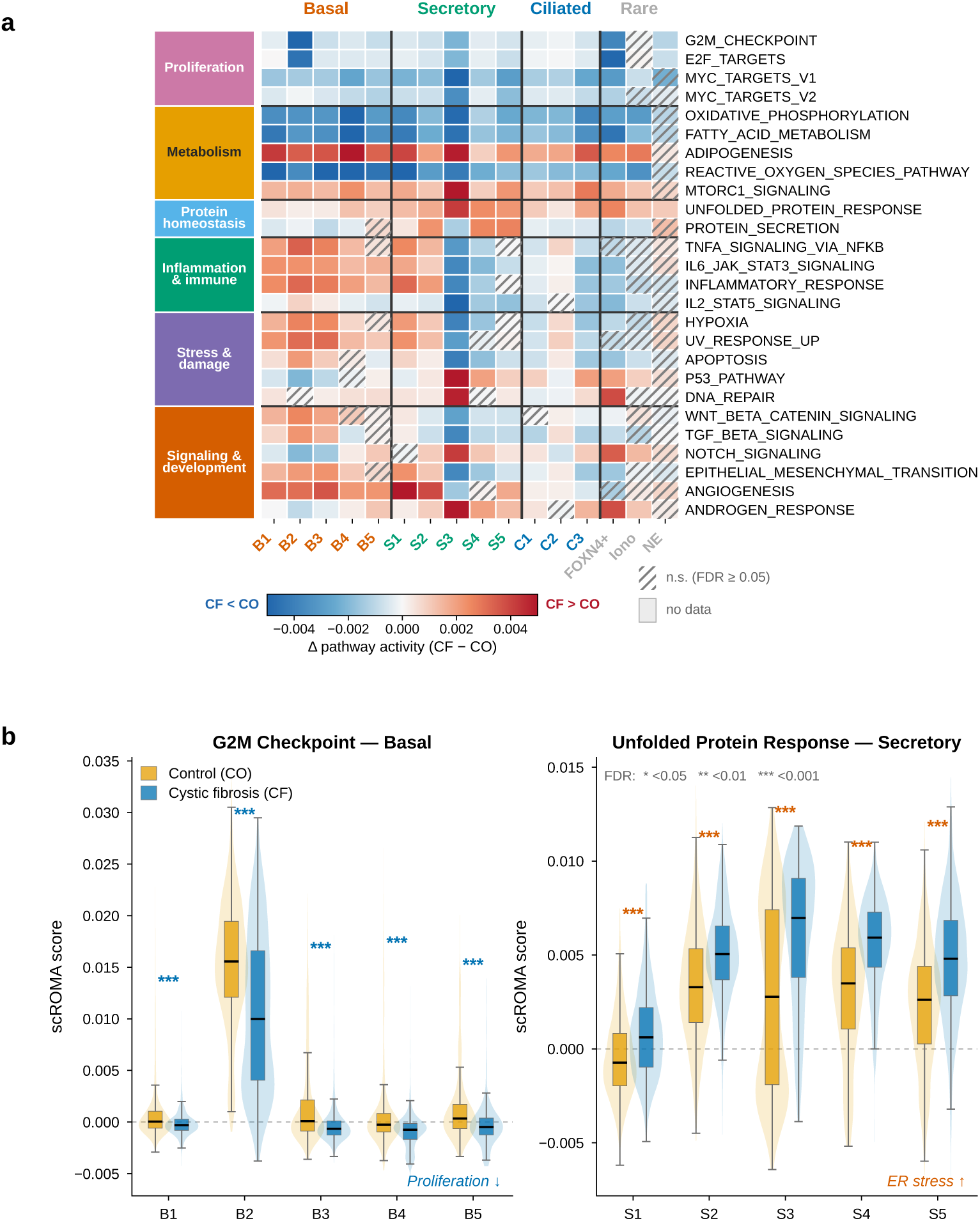
scROMA resolves cell-type-specific pathway activity alterations in cystic fibrosis airway epithelium. **(a)** Delta heatmap showing the CF-minus-CO difference in median pathway activity score for each of the 16 epithelial subtypes, across 26 significantly active Hallmark pathways [8] in 40,709 cells from CF (*n* = 19) and control (CO, *n* = 19) proximal airway epithelium [31]. Red indicates higher activity in CF; blue indicates higher activity in CO. Columns are ordered by cell-type hierarchy (Basal, Secretory, Ciliated, Rare); rows are organized into six functional pathway groups (color sidebar, left): proliferation, metabolism, protein homeostasis, inflammation and immune, stress and damage, and signaling and development. Hatched cells are non-significant (FDR ≥ 0.05) and gray cells indicate subtypes with insufficient cells; significance was assessed by two-sided Mann-Whitney *U* with Benjamini-Hochberg correction across 416 comparisons. **(b)** Violin and box plots comparing CF (blue) and CO (orange) per-cell pathway activity distributions for two findings concordant with Carraro et al. [31]. Left: G2M_CHECKPOINT across basal subtypes, showing reduced proliferative activity in CF, most prominently in the cycling Basal2 subset (Δ = -5.6 × 10^-3^, *p*_ad__j_ < 10^-41^, *|r|* = 0.37). Right: UNFOLDED_PROTEIN_RESPONSE across secretory subtypes, showing elevated ER stress in CF. Significance stars (* FDR < 0.05, ** < 0.01, * * * < 0.001) are colored by direction of change in CF (blue, decreased; orange, increased). Data were batch-integrated using scANVI (institutional batch covariate) prior to scROMA base-mode scoring. All statistical tests: two-sided Mann-Whitney *U* with Benjamini-Hochberg correction.

Pathway activity was organized primarily by cell identity: the three major epithelial lineages formed distinct programs, and CF and control cells of the same subtype consistently co-clustered, indicating that cell type — not disease status — is the dominant axis of variation, with disease effects superimposed as subtype-specific modulations (Supplementary Fig. S5). We therefore quantified disease-associated alterations within each subtype, computing the CF-minus-CO difference in median pathway activity (Mann-Whitney *U* on cell-level scores, Benjamini-Hochberg-corrected across 416 comparisons); 356 (85.6%) were significant at *p*_adj_ < 0.05 (Fig. 4a). The delta heatmap revealed a structured, non-uniform pattern: proliferative pathways were reduced in CF (concentrated in basal subtypes), oxidative phosphorylation and fatty-acid metabolism were reduced more broadly, and inflammatory and stress pathways shifted in a celltype-dependent manner — elevated in basal and early secretory cells, attenuated in ciliated and late secretory cells.

Several alterations directly recapitulated the key conclusions of Carraro et al. [31], despite scROMA using only generic MSigDB Hallmark gene sets rather than the custom co-expression networks derived in that work. Foremost was the proliferation deficit in CF basal progenitors, which they validated by PCNA/KRT5 immunostaining: scROMA found G2M_CHECKPOINT significantly reduced in CF across all five basal subtypes, strongest in the cycling Basal2 subset (Δ = -5.6 × 10^-3^, *p*_adj_ < 10^-41^, |*r*| = 0.37; Fig. 4b, left), with E2F—TARGETS concordant (*|r|* = 0.40 in Basal2). This supports their stem-cell-exhaustion hypothesis, in which prolonged epithelial turnover in the chronically inflamed CF airway depletes the proliferative capacity of basal progenitors. A second concordance was elevated UNFOLDED _PROTEIN_RESPONSE in CF secretory cells (Fig. 4b, right; strongest in Secretory4, *|r|* = 0.46, and Secretory2, |*r*| = 0.36), matching the ER-stress network of Carraro et al. and consistent with the increased secretory burden of the CF airway. Further subtype-specific concordances each matched a reported network: elevated NOTCH-SIGNALING in secretory cells (their Notch network), expanded EPITHELIAL—MESENCHYMAL—TRANSITION in Basal4 (enhanced basal-to-ciliated transition), and elevated APOPTOSIS in Basal2 (stem-cell exhaustion). TNFA_SIGNALING_VIA_NFKB was elevated in the early secretory subtypes (Secretory1 and Secretory2) but reduced in Secretory3 and Secretory4.

By contrast, IL6_JAK_STAT3_SIGNALING and INFLAMMATORY—RESPONSE showed mixed directionality — elevated in CF basal and early secretory cells but attenuated in ciliated and late secretory cells. This pattern’s interpretation warrants caution, as it may partly reflect the difference between scROMA’s coordinated gene-set scoring and the gene-by-gene network analysis of the original study (see Discussion).

In summary, scROMA applied to a complex, multi-institutional atlas using only standard Hallmark gene sets, independently recovered the major cell-type-specific pathway alterations that Carraro et al. identified through extensive custom network analysis — validating its biological reliability at atlas scale, while the cell-type-dependent inflammatory patterns illustrate its capacity to resolve pathway-level structure that complements gene-level analyses.

### 2.5 scROMA resolves cell-type-specific pathway activity in a human intestinal-organoid atlas

Single-cell RNA-seq of intestinal organoids is notoriously difficult to read at the pathway level: organoid cultures are heterogeneous, carry strong donor and batch effects, and yield less consistent signal than primary tissue, making cell-type- and perturbation-resolved biology hard to recover and to compare across experiments. We asked whether scROMA — which assigns every cell an interpretable, per-pathway activity score computed independently within each cell type — can extract reproducible, cell-type-resolved pathway biology from such data. We applied it to the Capeling et al. human intestinal-organoid perturbation atlas [40] (144,146 epithelial cells passing quality control; three donors; 79 individual secreted niche factors plus an inflammatory cytokine cocktail [“cytomix”: TNF-*α* + IFN-*γ* + IL-1*β*] and a bare-medium control). scROMA scores were computed independently within each of four epithelial compartments — stem, transit-amplifying (TA), colonocyte, and goblet (136,842 cells across the four) — and each pathway’s latent axis was oriented to its member-gene expression independently of the cytomix label, so that orientation does not bias the recovery analysis. Rather than applying scROMA’s batch-aware correction with donor as the batch variable, which would remove the between-donor variance before it is used to test reproducibility, we scored each donor independently and required consistency across all three, an independent-replication test of recovery. We then asked (i) whether the score recovers a known perturbation and (ii) whether it resolves cell-type-specific responses to individual ligands.

#### Recovery of the inflammatory program across cell types (Fig. 5a)

For each pathway, cell type, and donor, we measured how well the scROMA score separates cytomix-treated (inflamed) from bare-medium (homeostatic) cells, summarized as the area under the ROC curve (AUC; 0.5 = no separation, 1.0 = perfect separation; >0.5 = the pathway is induced by cytomix). scROMA recovered the cytomix inflammatory program in all four cell types and reproducibly across all three donors: the KEGG inflammatory-bowel-disease signature [38] (AUC 0.87−0.92), the interferon-*α* and -*γ* responses (0.71−0.95), interferon-*α*/*β* signaling (0.79−0.91), IL6-JAK-STAT3 signaling (0.69−0.87), and allograft rejection (0.82−0.94). The signal was programspecific rather than a global increase in pathway activity: unrelated metabolic and structural pathways stayed at or below chance (oxidative phosphorylation 0.41−0.48, fatty-acid metabolism 0.42−0.52, Notch 0.51−0.53), a built-in specificity control. The magnitude of recovery was cell-type-graded — the interferon-*γ* response was strongest in TA and colonocytes (0.92) and attenuated in goblet cells (0.71), while IL6-JAK-STAT3 peaked in stem cells (0.87) — highlighting that the same stimulus engages each compartment to a different degree.

**Fig. 5:**
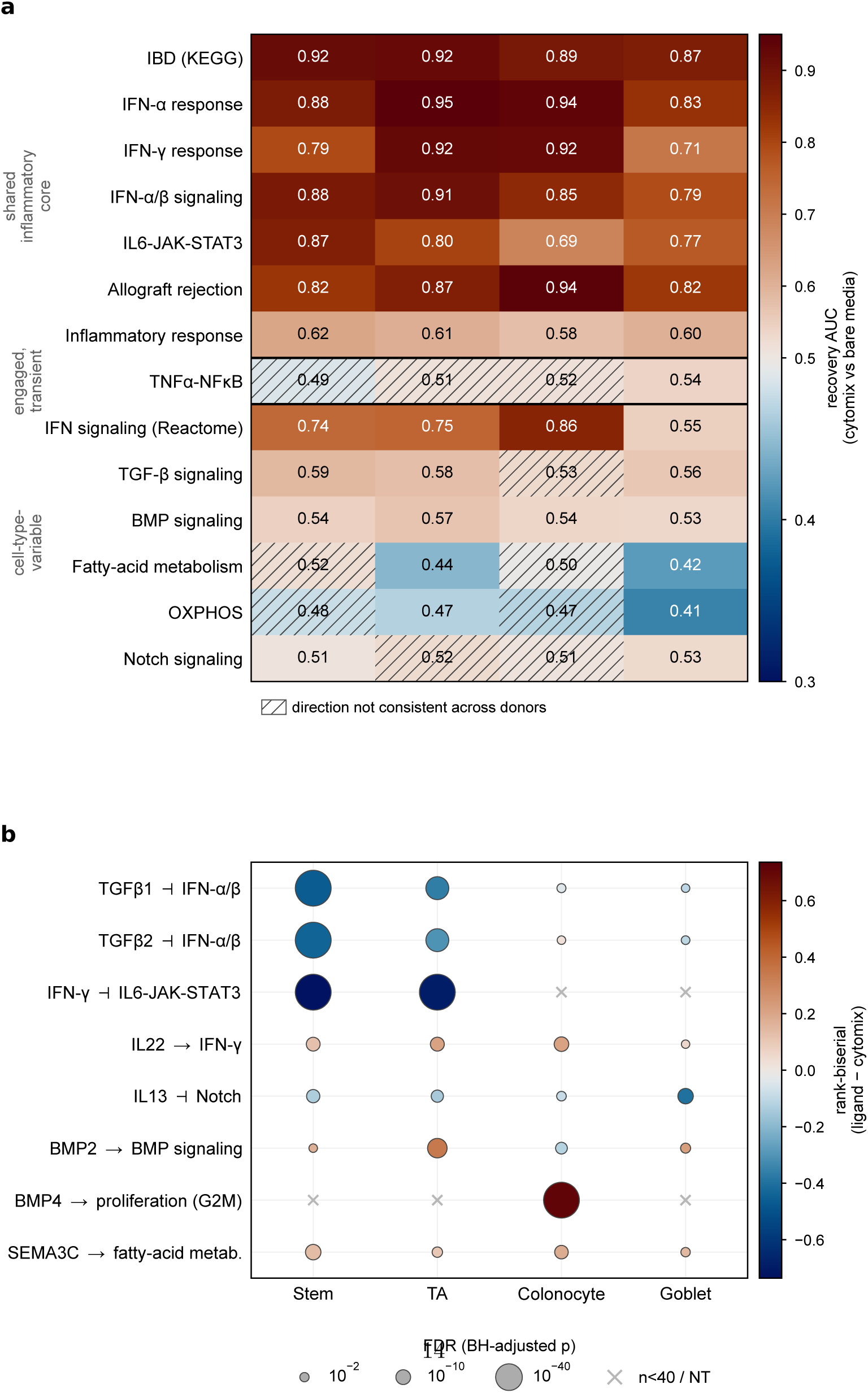
scROMA recovers a cell-type-resolved inflammatory program and resolves cell-type-specific ligand targets in a human intestinal-organoid perturbation atlas. scROMA pathway-activity scores computed on the Capeling et al. atlas [40] (144,146 epithelial cells; three donors; 79 secreted niche factors plus cytomix and bare medium), independently within each of four epithelial compartments; both panels re-use the same per-cell scores. **(a)** Recovery heatmap. For each pathway × cell type, the color and printed value give the ROC-AUC (mean of three donors) of the single-pathway score separating cytomix from bare-medium cells (AUC 0.5 = no separation, 1.0 = perfect; >0.5 = induced by cytomix). Rows are grouped into a shared inflammatory core recovered in every compartment; TNF-a-NF-*κ*B shown separately as “engaged but transient”; and cell-type-variable programs, including the unrelated metabolic/structural specificity controls (oxidative phosphorylation, fatty-acid metabolism, Notch). Hatched cells mark pathway × cell-type combinations whose effect direction is not consistent across all three donors. **(b)** Ligand-target selectivity dot plot. For curated ligands, the rank-biserial effect on the ligand’s signature pathway relative to the cytomix baseline, pooled across donors per compartment (color, -1 to +1; ˧ = suppresses, → = activates); dot size encodes the Benjamini-Hochberg-adjusted significance (-log_10_ FDR); “×” marks combinations with too few cells to test (*n* < 40 / not tested). Both panels share the same four-compartment x-axis (Stem, TA, Colonocyte, Goblet).

One inflammatory module behaved differently and is shown as its own category (“engaged but transient”): TNF-α-NF-*κ*B stayed at chance (AUC 0.49−0.54) even though TNF-*α* and IL-1*β* — both NF-*κ*B agonists — are components of cytomix. This reflects timing rather than a method failure: NF-*κ*B is an acute, selhlimiting program (its target transcription peaks within a few hours and is auto-terminated by IκB*α*/A20), whereas the interferon and JAK-STAT responses are sustained, and the atlas is read out well after the cytomix pulse. A raw, scROMA-independent geneexpression analysis confirms this interpretation (Supplementary Fig. S6): the NF-*κ*B Hallmark module barely separates cytomix from bare medium (mean-expression AUC 0.53−0.55) while the interferon-*γ* module separates strongly (0.88−0.97), and the fastest immediate-early NF-*κ*B autoregulators (NFKBIA, IER3, NFKB1) have returned to baseline. scROMA thus reads the temporal structure of the response — capturing the programs still active at readout — rather than reporting every cytokine that was added.

#### Cell-type-specific ligand targets (Fig. 5b)

For individual ligands, we computed the rank-biserial effect size (-1 to +1; sign relative to the inflamed cytomix baseline, negative = suppression [˧], positive = further activation [→]; |effect| ≈ strength) on each ligand’s signature pathway, per cell type. scROMA pinpointed targets that are sharply cell-type-restricted. TGF-*β*1 and TGF-e2 suppressed interferon-*α*/*β* signaling selectively in stem (−0.46/−0.44) and TA (−0.36/−0.31) cells, with no significant effect in colonocytes or goblet cells. IFN-*γ* markedly lowered IL6-JAK-STAT3 activity relative to the cytomix background in stem (−0.74) and TA (−0.69) cells (untestable in colonocyte and goblet cells, too few ligand cells). IL22 mildly raised the interferon-*γ* program in stem, TA, and colonocyte cells (+0.12 to +0.21). IL13 suppressed Notch most strongly in goblet cells (−0.40). BMP2 engaged BMP signaling most clearly in TA cells (+0.34), and BMP4 drove a strong proliferative (G2M) program specifically in colonocytes (+0.71). Finally, SEMA3C — an under-characterized ligand not emphasized in the original study — up-regulated fatty-acid metabolism, most strongly in colonocytes (+0.18), nominating a concrete, testable hypothesis for follow-up. Where a ligand had too few cells in a compartment to test, the effect is not reported (marked “×”).

Together, these results show that scROMA reads pathway activity at single-cell, cell-type resolution in a challenging organoid setting: it sensitively and reproducibly recovers a known inflammatory signal in every epithelial compartment and across donors, distinguishes the temporal structure of that response (sustained interferon versus transient NF-*κ*B), and resolves the cell-type-specific pathway targets of individual ligands — capabilities that matter precisely because organoid single-cell data are heterogeneous and hard to compare across experiments.

### 2.6 Batch-aware scROMA resolves subtype-specific and cell-type-specific pathway activities in breast cancer

The breast cancer tumor microenvironment is remarkably heterogeneous, comprising cancer epithelial cells spanning molecularly distinct intrinsic subtypes, diverse immune populations, and a complex stromal milieu — all shaped by the clinical subtype of the disease (ER^+^, HER2^+^, TNBC) [32, 39]. Resolving this at the pathway level requires separating patient-specific technical variation from subtype- and cell-type-specific biology. We therefore applied scROMA in batch-aware mode, with patient identity as the batch variable, to a breast cancer scRNA-seq atlas [32] of 26 primary tumors — ER^+^ (*n* = 11), HER2^+^ (*n* = 5), and TNBC (*n* = 10) (see Methods; Fig. 6).

**Fig. 6:**
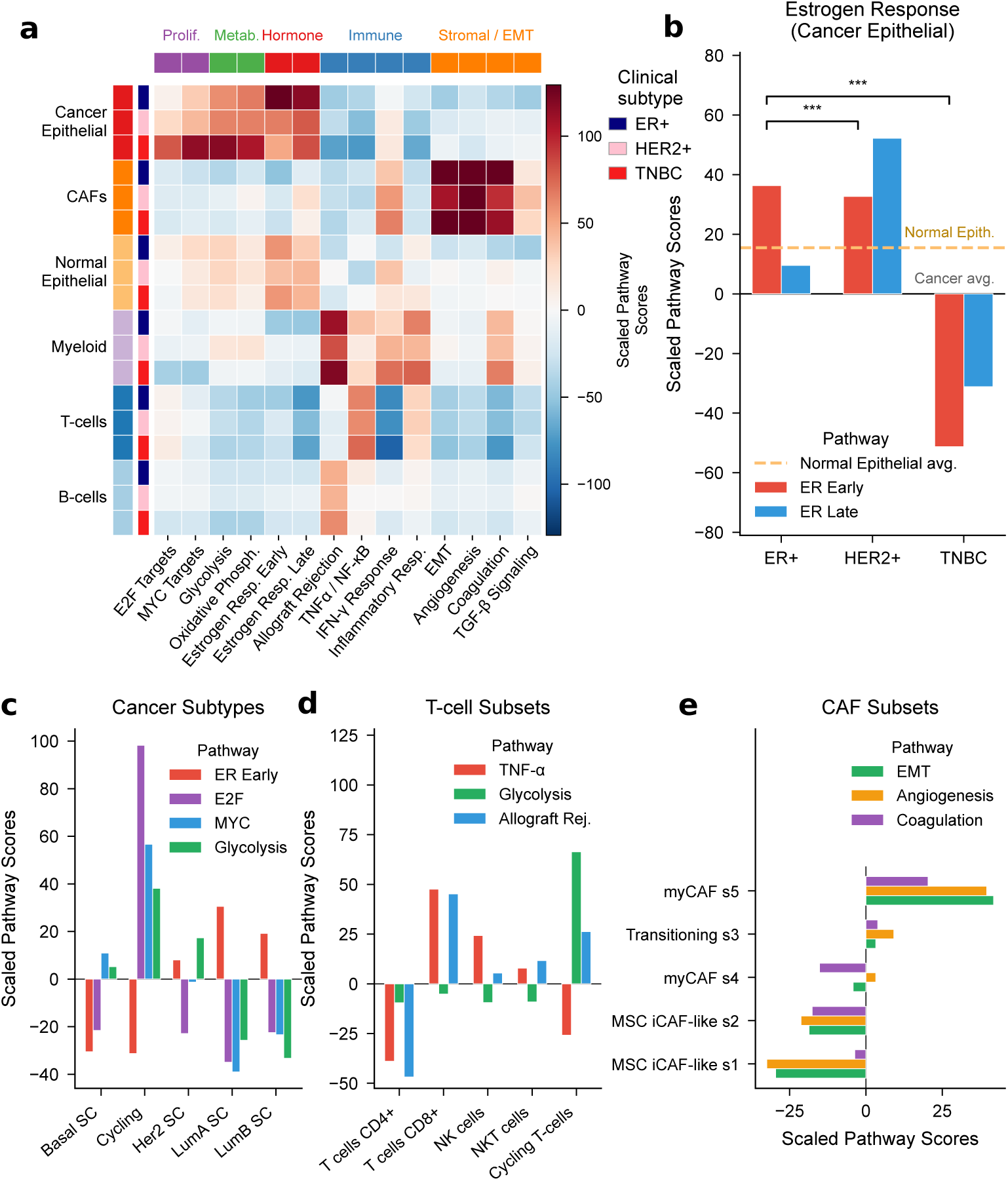
Batch-aware scROMA resolves multi-level pathway activity in the breast cancer tumor microenvironment. **(a)** Pathway activity heatmap across ma jor cell-type compartments and clinical subtypes (ER**^+^**, HER2**^+^**, TNBC) in 26 primary breast tumors [32], scored by scROMA in batch-aware mode (patient as batch variable). **(b)** Estrogen-response activity (early and late) in cancer epithelial cells by clinical subtype, scored relative to the cancer-epithelial average (zero line); the dashed line marks the mean of the two per-pathway normal-epithelial values on the same scale. ER**^+^** and HER2**^+^** are estrogen-active and TNBC negative (P < 0.001). **(c)** Cancer-cell pathway profiles by SCSubtype (Basal, HER2-enriched, Luminal A, Luminal B, Cycling), concordant with independently derived molecular subtypes. **(d)** T-cell and innate lymphoid subset pathway profiles (TNF-a/NF-*κ*B, allograft rejection, glycolysis), resolving functional specialization of CD8**^+^**, CD4**^+^**, NK, NKT-like, and cycling T-cells. **(e)** CAF subset profiles (EMT, angiogenesis, coagulation) along the MSC/iCAF-to-myCAF trajectory.

#### Global pathway landscape (Fig. 6a)

Across major cell-type compartments and clinical subtypes, scROMA recovered a strongly structured landscape consistent with established breast cancer biology. Cancer epithelial cells were enriched for proliferation (E2F and MYC targets), metabolic (glycolysis, oxidative phosphorylation), and hormone-response pathways; cancer-associated fibroblasts (CAFs) for stromal and EMT programs (EMT, angiogenesis, coagulation, TGF-*β* signaling); and the immune compartments (T cells, B cells, myeloid cells) for immune pathways (allograft rejection, TNF-*α*/NF-*κ*B, IFN-*γ* response, inflammatory response). This clean separation along cell-type and functional axes shows that batch-aware scoring preserves the expected biological structure despite the initial substantial inter-patient variability.

#### Estrogen response tracks clinical subtype (Fig. 6b)

As a positive control, estrogen-response activity in cancer epithelial cells followed clinical subtype: scored relative to the cancer-epithelial average, with the normal-epithelial level marked for reference, ER**^+^** and HER2**^+^** tumors were estrogen-active (early and late) whereas TNBC was strongly negative (*P* < 0.001), recapitulating the hormone-receptor biology that guides clinical treatment. HER2**^+^** cells showed unexpectedly high estrogen-late activity, exceeding ER**^+^** on average; the per-patient analysis shows this to be uniform rather than driven by outlier tumors — in all three HER2**^+^** tumors contributing cancer epithelial cells, 89-99% of those cells are estrogen-late-active, the tightest such range of any subtype, whereas ER**^+^** (37-100%) and TNBC (32–94%) are markedly more heterogeneous (Supplementary Fig. S7), consistent with the intratumoral subtype heterogeneity evident across patients (Supplementary Fig. S7c).

#### Cancer-cel l subtype profiles (Fig. 6c)

Within cancer epithelial cells, scROMA profiles aligned with the independently derived single-cell SCSubtype assignments of Wu et al. [32]: cycling cells had the highest E2F, MYC, and glycolysis activity; HER2-enriched cells elevated glycolysis; and the luminal subtypes (Luminal A, Luminal B) were the cancer subtypes with strong estrogen-response-early activity while being the most strongly depleted for E2F and MYC — the hormone-driven, low-proliferative profile that defines luminal cancers clinically. This concordance with an orthogonal single-cell classification, and with the gene-module analysis of Wu et al. (a method-level comparison is given in the Discussion), validates scROMA’s capacity to capture biologically meaningful pathway variation at single-cell resolution.

#### T-cell subset specialization (Fig. 6d)

Among lymphoid subsets, CD8_+_ T cells had the highest TNF-α/NF-*κ*B and allograft-rejection activity, the latter gene set being enriched for granzyme and perforin effectors, consistent with their cytotoxic role, with NK and NKT-like cells following for TNF-a/NF-ΚB and cycling T cells second for allograft rejection. Cycling T cells were distinct: highest glycolysis and strong effector activity but markedly negative TNF-α signaling, reflecting the metabolic shift of proliferating T cells coupled with suppressed NF-ΚB signaling, while CD4**^+^** cells scored low across these effector pathways. These distinctions match the lymphoid populations Wu et al. [32] described.

#### CAF state specialization (Fig. 6e)

Across the five CAF states of Wu et al. [32], stromal activity formed a gradient along the iCAF-to-myCAF trajectory: the ECM-producing myofibroblastic state myCAF s5 had the highest EMT, angiogenesis, and coagulation activity; the transitioning s3 state ranked second, consistent with its mixed iCAF/myofibroblast identity; and the inflammatory MSC/iCAF-like s1 and s2 states the lowest. CAFs in TNBC tumors showed higher IFN-*γ* response activity than those in ER_+_ or HER2_+_ tumors (Fig. 6a), consistent with the inflamed TNBC microenvironment. These fine-grained distinctions illustrate scROMA’s resolution of functional heterogeneity within a single compartment.

Taken together, these results demonstrate that batch-aware scROMA resolves the breast cancer pathway landscape at multiple levels: clinical-subtype distinctions in cancer epithelial cells, functional specialization of immune subpopulations, and stromal remodeling states in fibroblasts — consistent with the biology described by Wu et al. [32] and recovered despite substantial inter-patient batch effects.

### 2.7 Pathway activity scores capture copy number variation-associated functional heterogeneity in lung cancer

Beyond scoring individual pathways, we asked whether the full scROMA profile could serve as a biologically interpretable embedding that reflects tumor genomic architecture. We applied scROMA (base) to a lung cancer scRNA-seq dataset (Maynard et al. [33], *n* = 3,000 cells), scoring all 50 Hallmark gene sets [8], of which 43 were significantly shifted by scROMA’s permutation test (FDR < 0.05). We then compared clustering in this 43-dimensional pathway embedding against two other pathway-scoring methods applied to the same pathways — AUCell [11] and Scanpy’s `score—genes` [13] — and against gene-expression PCA (50 components), using inferred clonal structure as the reference (Fig. 7).

**Fig. 7:**
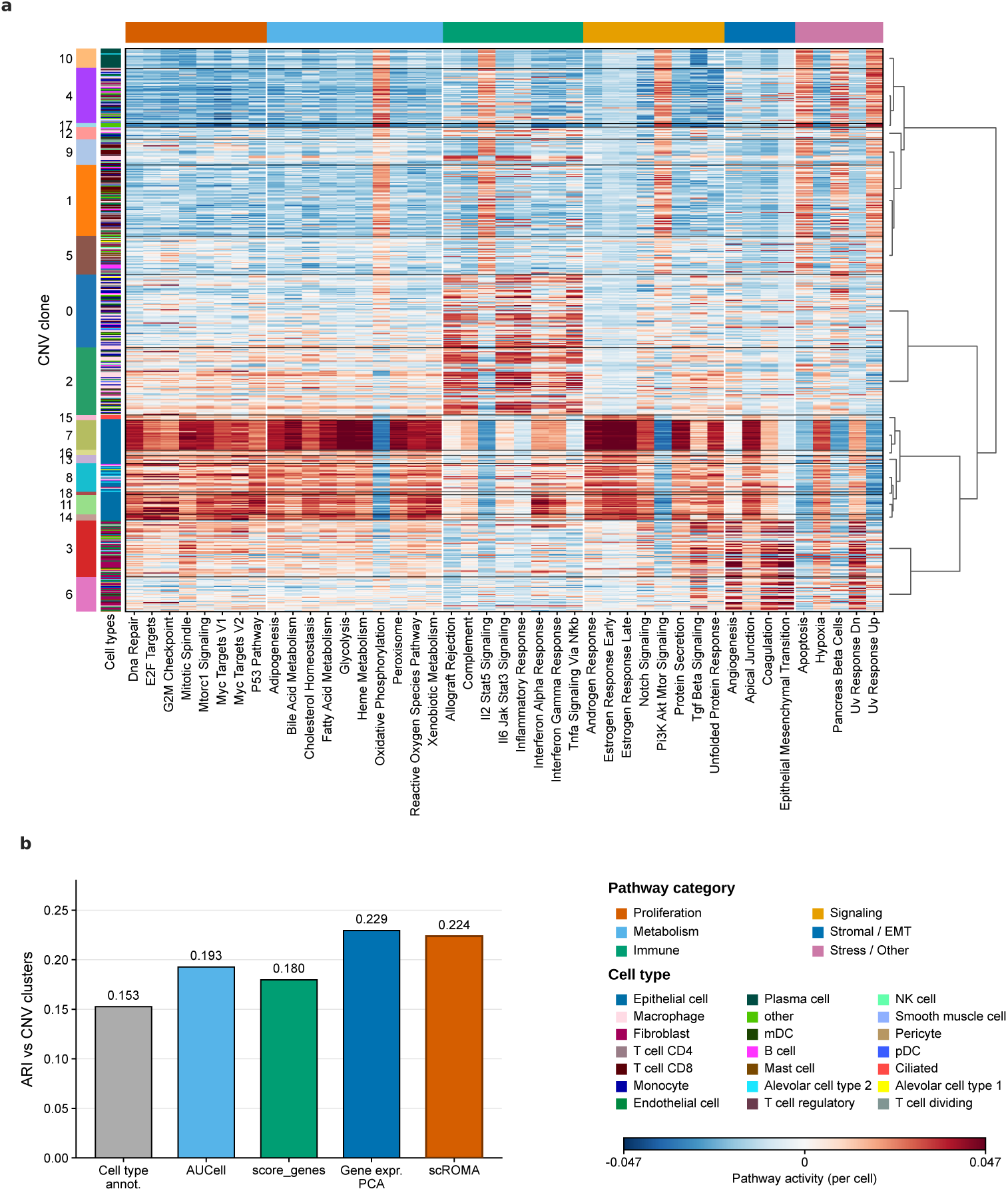
scROMA pathway activity scores capture CNV-associated functional heterogeneity in lung cancer. **(a)** Heatmap of scROMA per-cell pathway activity scores in the lung cancer dataset [33]. Cells are grouped by CNV-derived clonal cluster (rows; left annotations give clone identity and cell-type composition), and the 43 significantly active Hallmark pathways (columns) are grouped by functional category (top bar). Distinct CNV clones show characteristic pathway profiles not explained by cell-type composition; the dendrogram (right) reflects hierarchical clustering of clones by pathway activity. **(b)** Adjusted Rand index (ARI) between clustering in each representation and the CNV clonal clusters, for cell-type annotation, AUCell, `score_genes`, gene-expression PCA, and scROMA. scROMA (0.224) matches gene-expression PCA (0.229) and exceeds the other pathway-scoring methods, while all expression-based embeddings surpass cell-type annotation (0.153).

We inferred copy-number variation (CNV) profiles from expression with inferCNVpy [42, 43], and defined clonal clusters by Leiden clustering in CNV space (see Methods); the highest-CNV clones corresponded to the epithelial (tumor) compartment (Supplementary Fig. S8). These CNV clones aligned only weakly with transcriptomic cell-type identity (adjusted Rand index, ARI = 0.153), confirming that clonal structure and cell type are largely orthogonal axes of tumor heterogeneity. Consistent with this, the scROMA pathway-activity heatmap resolved the clones into groups with characteristic pathway profiles — spanning proliferative, metabolic, immune, and stromal programs — that were not explained by cell-type composition (Fig. 7a).

We then quantified how well clustering in each representation recovered the CNV clones (Fig. 7B). All expression-based embeddings outperformed cell-type annotation alone (ARI = 0.153), indicating that pathway activity carries clonal-genomic information beyond cell-type identity. Critically, scROMA’s interpretable pathway embedding (ARI = 0.224) matched gene-expression PCA (0.229) — the strongest baseline, which operates on thousands of raw genes rather than 43 labeled pathways — and clearly exceeded the other pathway-scoring methods (AUCell 0.193, `score_genes` 0.180).

All panels show the standardized deviation of each group mean from a reference population, 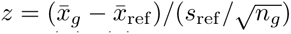. Panel **(a)** uses all cells as the reference population; panels **(b)—(e)** use within-cell-type references. Magnitudes are therefore comparable within a panel but not across panels with different reference sets. scROMA thus compresses the transcriptome into a compact, biologically labeled representation that captures clonal genomic structure as faithfully as an uninterpretable gene-level embedding.

Together, these results show that scROMA pathway scores form an interpretable embedding in which clonal genomic structure is recoverable, positioning pathway activity as a bridge between transcriptomic cell state and genomic architecture.

## 3 Discussion

Taken together, these analyses show that pathway activity in multi-sample single-cell data can be made statistically grounded and batch-aware within a single scoring step, recovering biology that whole-transcriptome integration and gene-by-gene testing each leave partly hidden.

### Correcting batch effects inside each gene set

scROMA’s central choice is to remove batch effects within a pathway’s own genes rather than across the full expression matrix. Whole-transcriptome integration is tuned to blend cells globally and can flatten the pathway-level differences a score is meant to measure; correcting inside the gene set removes only the technical variation among that pathway’s genes and leaves the rest of the biology intact. This correction should be applied only when batch is a nuisance, not the effect of interest: because it erases variation along whatever axis it is given, we scored the organoid atlas in base mode — where the aim was to test reproducibility across donors and used batch-aware mode only for the breast atlas, where between-patient variation is genuine technical noise.

### Coordinated scoring is complementary to gene- and module-level analysis

Summarizing a gene set by its dominant coordinated axis interrogates a different layer of regulation than differential expression, which reconciles both the agreements and divergences with prior work. Concordance was strongest for broad, tightly coordinated programs — proliferation and ER stress in the cystic fibrosis epithelium [31] — where a coordinated summary and a gene-level analysis should agree. Divergence, such as the cell-type-dependent inflammatory signal there, need not mean error in either: a pathway can shift coherently in its dominant mode when many genes change by amounts too small to pass a per-gene threshold, while a few strongly regulated genes need not move that mode. The same mechanism is an asset when programs are bundled — scoring each Hallmark set independently separates proliferation from glycolysis in cancer subtypes that composite modules score as one [32]. Coordinated scoring thus reports a program’s aggregate functional state, best read alongside gene-level results.

### The score reflects the present biology

A permutation-anchored SVD score tracks the transcriptional state actually realized in the profiled cells. In the organoid atlas it reported the sustained arm of an inflammatory stimulus but not its already-resolved acute NF-*κ*B component — reading the response’s temporal structure rather than the cytokines administered, as an orthogonal raw-expression analysis confirmed. In the lung dataset, the same permutation-anchored profile acts as a compact, biologically labeled embedding that tracks clonal genomic structure as well as an uninterpretable gene-level one (Fig. 7; Supplementary Fig. S9; [33]).

### A benchmark with exact ground truth

The simulation framework’s value lies less in any single result than in what it enables: pathway methods can be compared against activation states that are known rather than inferred. Its deliberate simplicity — a transparent generative model in place of maximal realism — makes each method’s behavior attributable to a defined cause, and it was this design that exposed the ranking collapse under batch effects that no single accuracy metric revealed. Released with pre-generated datasets [49], it is meant to serve as a pathway benchmarking for a wider scientific community.

### Implications for organoid systems

Organoid single-cell data are a demanding case for functional interpretation: cultures are heterogeneous, carry strong donor and batch effects, and give less reproducible signal than primary tissue, which makes cell-type- and perturbation-resolved biology hard to recover and, above all, hard to compare across experiments. A coordinated, per-cell, batch-aware pathway score suits these conditions — more robust to singlegene noise than gene-level readouts, equipped with a significance test that separates engaged programs from background, and, as a compact and biologically labeled summary, able to place perturbations, donors, and studies in a common coordinate system. For the niche-factor and perturbation atlases now being generated, such a representation offers a route to quantitatively comparable, cell-type-resolved pathway responses [40] that should extend well beyond the intestinal system examined here.

### Limitations

Like all SVD-based scores, scROMA summarizes each gene set along a single axis: overdispersion and cell-level rankings are sign-invariant, but the direction of a shift depends on orienting the leading singular vector, which we fix by anchoring to member-gene expression and corroborating against independent signatures [14] — principled, but an assumption, and one a single component cannot extend to pathways with several regulatory axes. The batch-aware strategies are largely linear or iterative-linear and will not fully resolve strongly non-linear confounding, and, as above, subspace correction can remove real biology when the batch label is entangled with the contrast of interest. The simulator’s log-linear Poisson model omits negative-binomial overdispersion, zero-inflation beyond expression-dependent dropout, and gene-gene correlation beyond that imposed by pathway membership. Finally, permutation testing and, in batch-aware mode, leave-one-out filtering dominates runtime at very large gene-set scale, where the prototype sparse operator and analytical null approximations are the clearest routes to genome-scale performance. Extending subspace correction to nonlinear methods, exposing multi-component representations, and applying scROMA to spatial and multi-modal data are the natural next steps.

## 4 Conclusions

scROMA moves batch correction to the level of the gene set, couples SVD pathway scoring with permutation significance and per-cell resolution in the Scanpy/AnnData ecosystem [13], and ships with a ground-truth simulation framework the field has lacked. As open-source software with pre-generated benchmarks and reproducible analysis scripts, it is intended to make statistically grounded, batch-aware pathway analysis a routine step in single-cell workflows.

## 5 Methods

scROMA is an open-source Python package that operates natively on AnnData objects and stores all results within the AnnData structure, so that pathway scores integrate directly into downstream Scanpy [13] workflows (clustering, embedding, and differential testing). The subsections below describe, in the order of the Results, the core algorithm (5.1), the batch-aware extension (5.2), the ground-truth simulation framework (5.3), the benchmarking and evaluation design (5.4), and the five real-data analyses (5.5).

### 5.1 The scROMA algorithm

scROMA is a Python implementation of the ROMA (Representation and Quantification Of Module Activity) framework [29] for single-cell and bulk transcriptomics. It takes two inputs: a gene-expression matrix as an AnnData object (cells × genes) and a collection of gene sets in Gene Matrix Transposed (GMT) format, as distributed by MSigDB [8], KEGG [38], Reactome, or any user-defined dictionary. The expression matrix is expected to be library-size-normalized and log-transformed: the same preprocessing applied before principal-component analysis, but *not* scaled to unit variance per gene, since such scaling would suppress the expression-magnitude differences that the overdispersion and gene-contribution scores depend on. Gene sets with fewer than a minimum number of member genes present in the matrix (default 10) are not scored. For each remaining gene set, scROMA returns two complementary pathway-level statistics—the L1 overdispersion score and the median-expression shift score—together with per-cell activity scores, per-gene contribution weights, and permutation-based significance values.

#### Centering

The expression matrix is transposed to a genes × cells orientation and centered prior to decomposition. By default scROMA applies sequential centering: each gene is centered across cells, and the resulting matrix is then centered across genes, removing both gene-level baseline expression and per-cell offsets from the subspace. An exact doublemean-centering mode 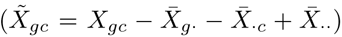, matching the original ROMA formulation, is available as an option. Centering statistics are computed once on the full matrix, so every gene-set submatrix is decomposed in a common reference frame.

#### SVD-based pathway scoring

For a gene set *G_k_* with |*G_k_*| genes present in the data, scROMA extracts the centered submatrix 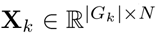 (genes × cells, N cells) and computes a truncated singular value decomposition (SVD), **X**_*k*_ ≈ **UƩV**^T^, where **Ʃ** = diag(σ_1_, σ_2_). Two components are retained internally (the second is used only to inspect the gap between the leading singular values); all scoring is based on the first. The decomposition is computed with scikit-learn’s randomized `Truncated-SVD` [35, 37], with an exact ARPACK solver available as an alternative. From the first component scROMA derives four quantities. The *L1 score* is the fraction of submatrix variance captured by the first principal component,

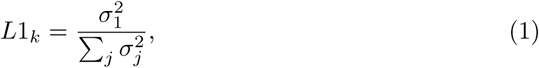

so that a high value indicates that expression among the pathway’s genes is concentrated along a single axis (overdispersion) more than for a random gene set of equal size. Gene-level projections onto the first principal direction, **p = X**_*k*_**v_1_** (with **v**_1_ the sign-oriented first right singular vector), give both the *median-expression score* MedExp*_k_* = median(**p**)—a signed measure of coherent up- or down-regulation (shift)—and the *per-gene contribution weights* **p**, whose magnitude and sign identify the genes driving the signal. The *per-cell activity scores* are the projections of individual cells onto **v**_1_, providing single-cell pathway activity. Overdispersion and shift are complementary: the former measures the magnitude of coordinated variability, the latter its direction and coherence.

#### Orientation of the first principal component

Because the sign of a singular vector is arbitrary, scROMA orients **v**_1_ per gene set so that positive scores correspond to pathway activation. Under the default `PreferActivation` mode, the genes whose absolute projection exceeds the *τ*-th quantile of all absolute projections (default *τ* = 0.90, i.e. the top-contributing decile) are identified, and the sign of **v**_1_ is flipped if the sum of their projections is negative. Additional modes use user-supplied gene weights or member-gene expression to orient the component when prior directional knowledge is available. `PreferActivation` governs orientation unless a per-analysis rule is specified; for the analyses whose direction is biologically interpreted (5.5), each pathway’s sign is instead anchored to member-gene expression, so that no directional claim rests on the arbitrary singular-vector sign. Orientation is applied independently for every gene set, so directional statements are not confounded across pathways.

#### Outlier-gene detection

To assess whether scores reflect coordinated expression rather than a few dominant genes, scROMA performs leave-one-out cross-validation (LOOCV) within each gene set: for each member gene, the single-component L1 score is recomputed on the remaining genes, and a gene is flagged when its leave-one-out z-score exceeds 3.0. A complementary cross-gene-set filter tracks, for every gene, how often it is flagged relative to how many gene sets contain it, and applies a one-sided Fisher’s exact test; genes with *p* < 0.05 that occur in at least three gene sets are excluded globally, preventing systematically variable genes (e.g. ribosomal or stress-response genes) from distorting multiple decompositions. To preserve small gene sets, at most 50% of a set’s genes may be removed, prioritized by LOOCV z-score. In batch-aware mode, flagged genes are removed before the final SVD. In base mode — the mode used for the cystic fibrosis, organoid and lung analyses — these *z*-scores are computed and returned as a diagnostic but no gene is removed, so every base-mode score reported here uses the complete set of member genes present in the data.

#### Significance testing

For each gene set of effective size *n*_k_ (after outlier removal), scROMA builds a null distribution from *N*_perm_ random gene sets of the same size, drawn from the full gene universe and processed through the identical pipeline (LOOCV filtering, SVD, scoring). The overdispersion p-value is one-sided,

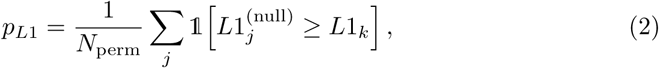

and the shift *p*-value is two-sided with a pseudocount,

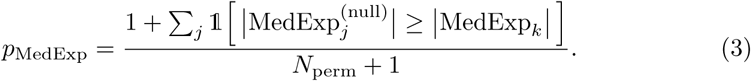

In batch-aware mode the pseudocount is applied to both statistics, so the smallest attainable *p*-value is 1/(*N*_perm_ + 1); this is the shared minimum q-value seen in Supplementary Table S3. Raw *p*-values are corrected across all tested gene sets by the Benjamini-Hochberg procedure [36]; a gene set is called significantly active when *q*_L1_ < 0.05 or *q*_MedExp_ < 0.05. Null distributions are cached by gene-set size and reused when effective sizes differ by no more than a tolerance (default 20 genes), and gene sets are processed in size order to maximize reuse.

#### Implementation and defaults

scROMA is implemented in Python 3 on NumPy, SciPy, scikit-learn, Scanpy, and AnnData, with null distributions computed in parallel via joblib. For memory-limited or atlas-scale settings, scROMA can decompose each submatrix without holding it densely in memory: an `IncrementalPCA` backend streams cells in mini-batches of user-defined size and closely approximates the truncated decomposition, and a sparse centered-SVD operator that represents the mean-centered submatrix implicitly—never materializing it—reduces scoring memory roughly four-fold in our benchmarks while reproducing the dense per-cell and pathway scores (Supplementary Note 2). The sparse operator is currently validated as a standalone prototype and is being integrated into the shipped interface; the results reported here use the default dense randomized `TruncatedSVD`. Compute time, memory, solver concordance, permutation detectability, batch-method cost, and CPU scaling are characterized in Supplementary Note 2.

Unless noted otherwise, analyses used LOOCV diagnostics with an outlier z-threshold of 3.0, a Fisher threshold of 0.05 (minimum three gene sets), a 50% per-set outlier cap, a null-size caching tolerance of 20 genes, randomized SVD, sequential centering, `PreferActivation` orientation with *τ* = 0.90, a significance threshold of q < 0.05, and a minimum gene-set size of 10. For permutation significance, a dedicated detectability analysis (Supplementary Note 2) identifies *N*_perm_ = 100 as the smallest null that reliably renders genuinely active pathways significant after multiple-testing correction on simulated ground truth, and this is the package default; because pathway effects in real data are typically subtler than in simulation and must be separated from a more heterogeneous background of random gene sets, we adopt a larger null of 300-700 permutations for the real-data applications, trading proportionally longer runtime for finer resolution of the smallest attainable p-values. Deviations are stated per analysis.

### 5.2 Batch-aware extension

To correct for batch effects scROMA applies correction within the gene-set subspace rather than to the full transcriptome. For each gene set *G_k_* it extracts the submatrix **X***_k_*, applies a batch correction driven by cell-level batch labels, and decomposes the corrected submatrix; because only the |*G_k_* | member genes are modified, variation encoded by genes outside the set cannot be removed. The identical correction is applied to every random gene set drawn for the permutation null (5.1), so significance is assessed against a null subjected to the same batch handling, and a dataset in which fewer than two batches are detected falls back to standard SVD.

scROMA provides seven correction strategies with a common interface (genes × cells submatrix plus a batch-label vector, returning a representation whose leading component feeds the standard scoring pipeline). *Centered* correction subtracts batch-specific gene means. *Residualized* correction regresses each gene on one-hot batch indicators by ordinary least squares (Ridge if *α* > 0) and retains the residuals. *Contrastive* PCA forms **Ʃ**_cPCA_ = **Ʃ**_total_ - *α* **Ʃ**_batch_ from the total and pooled within-batch covariances and projects onto the leading eigenvectors, maximizing biological variance while penalizing batch variance (*α* = 1.0 by default; a small diagonal shift restores positive-definiteness when needed). *Weighted* PCA downweights cells from high-variance batches by inverse batch variance. *ComBat* applies empirical-Bayes location/scale correction [22] within the subspace. *MNN* correction [34] estimates batch vectors from mutual nearest-neighbor pairs (default *k* = 20). *Harmony-like* correction [20] builds a low-dimensional cell embedding of the submatrix (default 20 PCs), soft-clusters it (default 50 clusters), and iteratively shifts batch-specific cluster centroids toward their global counterparts with batch-diversity reweighting (default 10 iterations; θ = 1.0). The first six operate in expression space and yield exact gene-level loadings; the Harmony-like variant operates in the embedding and derives equivalent gene contributions by projecting the original submatrix onto the corrected cell axis. When cell-type labels are supplied, the Harmony-like alignment is performed independently within each label, protecting known biological structure; without labels it runs unsupervised. Full mathematical definitions and default parameters for all seven strategies are given in Supplementary Methods.

### 5.3 Ground-truth simulation framework

To benchmark pathway-activity inference against exact ground truth, we developed a generative framework that produces synthetic scRNA-seq count matrices with fully specified per-cell pathway activities (Fig. 2a). It models count noise, expression-dependent dropout, library-size heterogeneity, and batch effects while retaining known activation states.

#### Gene universe and pathway architecture

We defined a universe of *G* = 1,000 genes and *P* = 50 pathways: 20 active (contributing structured signal) and 30 inactive decoys (annotated but contributing no coordinated signal). The active pathways span three architectures reflecting curated collections (Fig. 2d): a disjoint block (pathways 1-5; five non-overlapping sets of 25 genes) modeling cell-type markers; a high-overlap block (pathways 6-10; five sets of 40 genes drawn from a shared pool of 200) modeling functionally related programs; and a moderate-overlap block (pathways 11-20; ten sets of 20-50 genes from a pool of 400) reflecting the partial overlap typical of MSigDB and Reactome. Decoys (pathways 21-50; 15-60 genes) were sampled uniformly from the universe. All pathways were exported in GMT format.

#### Cell types and activation design

Five cell types were simulated with activation patterns spanning four scenarios (Fig. 2d): cell-type-specific (pathways 1-5, each active in one type), shared between pairs (pathways 6-10), globally active (pathways 11-13), and shared across combinations of two to three types (pathways 14-20). This combinatorial design yields ground truth for both per-cell detection and cell-type-level enrichment.

#### Ground-truth activities

Each cell type was assigned a monotonically decreasing sequence of mean activities across its active pathways, linearly spaced from *μ*_high_ = 2.0 to *μ*_low_ = 0.7, establishing a ground-truth ranking; non-associated pathways received *μ*_bg_ = 0.1. Per-cell activities were drawn as *s*_cp_ 〜 Ɲ(*μ*_t_(_c_)_,p_, *σ_s_*^2^) (truncated at zero, *σ_s_* = 0.3), and binary labels as 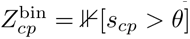 with *θ* = 0.5 (Fig. 2c).

#### Generative count model

Counts were drawn from a log-linear Poisson model. In the single-batch regime (*N* = 2,000 cells, equal cell-type proportions),

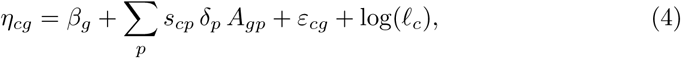

with baseline *βg* 〜Ɲ(-1.0, 0.6^2^), pathway effect *δ*_p_ 〜Ɲ(0.9, 0.2^2^), membership A_gp_ Ɛ {0, 1}, noise Ɛ*_cg_* 〜Ɲ(0, 0.7^2^), and library size *l*_c_ 〜 LogNormal(0.2, 0.5^2^). Counts were *X*_cg_ 〜Poisson(*λ*_cg_), *λ*_cg_ = exp(*η*_cg_) clipped to [10^-3^,100], with expression-dependent dropout *p*_drop_ = exp(-*λ*_cg_/*t*), *t* = 4.0. In the multi-batch regime (*B* = 3 batches of *N_b_* = 1,000 cells), the model added a global batch shift *γ_b_* 〜Ɲ(0,1.2^2^), gene-specific batch effects *B_bg_* 〜Ɲ(0, 0.8^2^), batch-dependent library shifts (*σ*_lib_=0.7), and per-batch cell-type compositions from Dirichlet(**1**). Pathway activities *s_cp_* were sampled identically across batches, so any drop in recovery reflects technical confounding rather than biological difference (Fig. 2b).

#### Replicates and output

Ten replicates per regime were generated with seeds 42—51 (identical parameters, distinct random draws), enabling mean±s.e.m. reporting. Datasets are released as AnnData objects carrying counts, cell-type and batch labels, continuous and binary ground-truth activity matrices, and all generative parameters [49]; the parameterized generator supports custom scenarios and records every generative parameter used.

### 5.4 Benchmarking and evaluation

#### Methods compared

We benchmarked scROMA against the three single-cell pathway-scoring methods, one per algorithmic family: AUCell [11] (ranking-based), GSEA [7] (enrichment-based), and Scanpy’s `score—genes` [13] (mean-expression, the Python equivalent of AddModuleScore [14]). AUCell and GSEA were run through decoupler [12] with default parameters (GSEA pruned to gene sets with at least two matched genes); `score—genes` used defaults (`ctrl_size`=50, `n_bins`=25). For scROMA we evaluated the base method and eight batch-aware arms: the seven correction strategies of 5.2, plus the Harmony-like strategy run without biological labels (`harmony_nobio`). The seven were passed the simulated cell-type annotation via `cell_type_key`; this affects only the Harmony-like strategy, whose centroid alignment is then performed independently within each label. The `harmony—nobio` arm withholds the annotation and is therefore the strictest, fully unsupervised setting, directly comparable to the scVI baselines, which likewise received only the batch key. In the multi-batch scenario we additionally benchmarked the conventional two-step workflow: scVI [21] was trained on the full count matrix (30 latent dimensions, 2 layers, negative-binomial likelihood, up to 300 epochs with early stopping), its corrected expression obtained via `get—normalized—expression`, and each competitor—plus scROMA base—applied to the scVI-corrected data (scVI+AUCell, scVI+GSEA, scVI+`score_genes`, scVI+scROMA).

#### Preprocessing and parameters

Simulated counts were library-size normalized and log-transformed; all 1,000 genes were retained so that gene sets stayed intact, and every method received the identical input (or the identical scVI-corrected matrix). scROMA was run with *N_p_*_erm_ = 300 for the benchmark; batch-aware variants used `batch_key.`=`“batch”` with the default parameters of 5.2. For GSEA, Task-2 rankings were taken from - log_10_ (q) rather than enrichment scores.

#### Evaluation tasks

Three tasks assessed methods at increasing granularity (Fig. 2e). Task 1 (*Activation Calling*) evaluated whether the 20 active versus 30 decoy pathways were correctly distinguished, and applies only to methods that report significance (scROMA, GSEA); AUCell and `score_genes` were marked not applicable. From the significance calls we computed the standard binary-classification metrics over the 20 active and 30 decoy pathways—sensitivity (recall of the active pathways, TP/(TP + FN)), specificity (TN/(TN + FP)), precision, the *F*_1_ score, balanced accuracy (the mean of sensitivity and specificity), and the Matthews correlation coefficient—and report sensitivity, specificity, and balanced accuracy (Fig. 3). *Task 2* (*Activity Ranking*) quantified preservation of the ground-truth pathway ordering by the absolute Spearman correlation between predicted and true rankings over the 20 active pathways. *Task 3 (Cell-Level Activity Recovery*) scored per-cell activity against the binary labels by AUC-ROC and AUC-PR and against activity by the absolute Pearson correlation, averaged over active pathways. All metrics are reported as mean±s.e.m. across the ten replicates (error bars in Fig. 3 denote s.e.m.).

#### Semi-controlled real-data benchmark

As a complementary benchmark with genuine biology, we augmented the Kang et al. PBMC IFN-*β* dataset [30] with a strong, fully specified synthetic batch effect and used the interferon response as ground truth (Supplementary Note 1). The full protocol, batch-effect parameters, and per-method results are given in Supplementary Methods.

### 5.5 Application to real datasets

Across all real-data analyses, pathway activity was scored with the MSigDB Hallmark collection v2023.1 [8] unless stated otherwise, cell-type annotations were taken from the original publications, expression was library-size normalized and log-transformed, and the random seed was fixed at 42.

#### Cystic fibrosis airway epithelium (Fig. 4)

We analyzed the Carraro et al. atlas [31] of proximal airway epithelium (GEO GSE150674): 40,709 cells from 19 cystic fibrosis (CF) and 19 control (CO) donors, spanning 16 epithelial subtypes processed at three institutions with distinct isolation protocols. To remove institution-level technical variation while preserving disease and cell-type structure, we integrated across the institutional covariate with scANVI [41] (institution as the batch key and the epithelial-subtype annotation as the cell-type label), then scored pathways with scROMA in *base* mode (institutional batch already addressed upstream) with *N*_perm_ = 1000, identifying 26 significantly active Hallmark pathways. We oriented every pathway’s PC1 so that positive scores correspond to higher mean log-normalized expression of its member genes. For the pathways carrying the principal directional findings we additionally corroborated this orientation against expression features independent of the score — a Tirosh cell-cycle signature for the proliferation programs, and curated marker panels for the oxidative-phosphorylation, fatty-acid-metabolism, reactive-oxygen-species, unfolded-protein-response, p53, and inflammatory programs across the basal, secretory, and ciliated lineages. Disease effects were quantified within subtype as the CF-CO difference in median activity, tested by two-sided Mann-Whitney *U* on per-cell scores with Benjamini-Hochberg correction across all 416 subtype × pathway comparisons; effect sizes are the rank-biserial correlation *r* = 1 - 2U /(*n*_CF_ *n*_CO_). The median-activity landscape (Fig. 4a) was hierarchically clustered over condition × subtype groups (average linkage, correlation distance; shown in Supplementary Fig. S5), and pathways were arranged into functional groups for display only.

#### Human intestinal-organoid niche-factor atlas (Fig. 5)

We analyzed the Capeling et al. atlas [40]: 144,146 epithelial cells from three donors exposed to 79 individual secreted niche factors, an inflammatory cytokine cocktail (“cytomix”; TNF-*α* + IFN-*γ* + IL-1*β*), and a bare-medium control. Scores were computed with scROMA in *base* mode independently within each of four epithelial compartments (stem, transit-amplifying, colonocyte, goblet; 136,842 cells), and each pathway was oriented to its member-gene expression independently of the cytomix label so that orientation does not bias the recovery analysis. Recovery of the inflammatory program was quantified per pathway, compartment, and donor as the area under the ROC curve separating cytomix-treated from bare-medium cells (0.5 = no separation, >0.5 = induced), summarized as the three-donor mean (Fig. 5a). Ligand-specific effects (Fig. 5b) were the rank-biserial effect size of each ligand on its signature pathway relative to the cytomix baseline, per compartment, pooled across donors, with Benjamini-Hochberg-adjusted significance; combinations with fewer than 40 cells were not tested.

#### Breast cancer tumor microenvironment (Fig. 6)

We analyzed the Wu et al. atlas [32] (GEO GSE176078): 26 treatment-naive primary tumors across ER**^+^** (*n* = 11), HER2**^+^** (*n* = 5), and TNBC (*n* = 10), with major, minor, and SCSubtype cell annotations from the original study. To separate patient-specific technical variation from subtype- and cell-type-specific biology, we scored pathways with scROMA in *batch-aware* mode using patient identity as the batch variable and the *Harmony-like* correction strategy (5.2) run without biological (cell-type) labels, so that inter-patient technical variation is removed without conditioning the correction on the cell-type and subtype structure that the analysis subsequently interrogates. The permutation null used the same correction, with *N*_perm_ = 700, and 27 Hallmark pathways were significantly active (Supplementary Table S3); the per-pathway outputs are deposited [50]. Summarizing each cell by its pathway scores yielded a patient-integrated embedding without a separate integration step, in contrast to unintegrated gene expression (Supplementary Fig. S10). The reported figures were computed on the full 100,064-cell atlas (after QC). For legibility, the landscape in Fig. 6a displays 14 representative pathways spanning the principal functional axes resolved across the microenvironment (proliferation, metabolism, hormone response, immune and stromal programs) rather than all 27, with the complete list of significantly active pathways provided in Supplementary Table S3; it shows, for each pathway and group g, the standardized deviation of the group mean from a reference population, 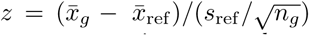; because the denominator carries 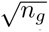, magnitudes grow with group size, so panels are comparable within, but not across, reference sets. Fig. 6a uses all cells as the reference population, whereas Fig. 6b-e use within-cell-type references. Estrogen response (Fig. 6b) was scored in cancer epithelial cells relative to the cancer-epithelial average, with the normal-epithelial level shown for reference; and the cancer-cell, T-cell and CAF subset profiles (Fig. 6c—e) apply the same statistic within each annotated state, pooled across patients.

#### Lung cancer copy-number heterogeneity (Fig. 7)

We analyzed the Maynard et al. lung adenocarcinoma dataset [33] (*n* = 3,000 cells across 21 annotated cell types), scoring all 50 Hallmark gene sets with scROMA in *base* mode; 43 were significantly shifted (FDR < 0.05) and defined a 43-dimensional per-cell pathway embedding. Copy-number profiles were inferred from expression with inferCNVpy [42, 43], taking the 11 immune cell types (B, plasma, NK, CD4**^+^**, CD8**^+^** and regulatory T, monocyte, macrophage, mast, mDC and pDC cells) as the normal reference and scoring the remaining epithelial, stromal and endothelial types against that pooled baseline, with a 250-gene running window at a 10-gene step, triangular window weighting, log-ratio clipping at ±3, a dynamic threshold of 1.5, and the sex chromosomes excluded (22 autosomes; 4,765 windows). The inferCNV-annotated 3,000-cell object on which these results were computed is deposited [50]. CNV clones were defined by Leiden clustering [44] of the CNV profile (50-component PCA, 15 nearest neighbors, Euclidean metric, resolution 1.0), yielding 19 clones that served as the reference partition. We then quantified how well unsupervised clustering in each of four cell representations—the 43-pathway scROMA embedding, AUCell and `score_genes` on the same 43 pathways, and gene-expression PCA (50 components)—recovered these clones, using the adjusted Rand index (ARI; `sklearn.metrics.adjusted_rand_score`), with cell-type annotation as a biological baseline. To keep the comparison fair, every representation used 15 nearest neighbors and was swept over an identical Leiden resolution grid (0.1—2.0) with the best-ARI solution reported; the pathway-score representations used a cosine metric suited to their scale, whereas gene-expression PCA used the Euclidean metric—the same metric and 50-component representation from which the reference clones were derived, an alignment that if anything favors the gene-expression baseline over scROMA. The gene-expression PCA and scROMA embeddings are visualized in Supplementary Fig. S9.

### 5.6 Software, data, and code availability

Analyses used scROMA (version 0.3.2) with Python 3, Scanpy [13], AnnData, scikit-learn [37], SciPy, and NumPy; scVI/scANVI via scvi-tools [21, 41]; AUCell and GSEA via decoupler [12]; CNV inference via inferCNVpy [42]; and matplotlib/seaborn for visualization. All random seeds were fixed (simulation replicates: 42—51; PBMC batch realizations: 0—5; all other analyses: 42). Public datasets: PBMC IFN-*β* (GEO GSE96583), CF airway (GSE150674), breast cancer (GSE176078), lung cancer (Maynard et al. [33]), and the intestinal-organoid atlas (Capeling et al. [40]). scROMA, the simulation framework, and the pre-generated benchmark datasets are released as open-source software with analysis scripts reproducing every figure; the software, the analysis code and all derived data are archived at Zenodo (see Availability of data and materials).

### 5.7 Use of large language models

A large language model (Claude, Anthropic; Sonnet and Opus model families, accessed 2025-2026) was used as an assistive tool during preparation of this work. Its use was limited to (i) language editing of manuscript text drafted by the authors, and (ii) refactoring, documentation and review of analysis and plotting code written by the authors. The model was not used to generate scientific content, to design or interpret analyses, or to produce results, figures or references. All model-assisted text and code were reviewed, verified and revised by the authors; all numerical results reported in this study were produced by the deposited scROMA code; and all cited references were verified against the primary literature. The authors take full responsibility for the content of this manuscript and for the correctness of the software.

## Supplementary information

Supplementary information accompanies this paper. It comprises Supplementary Methods, Supplementary Notes 1-3, Supplementary Figures S1-S10, and Supplementary Tables S1-S2.

## Abbreviations

ARI: adjusted Rand index
AUC: area under the curve
AUC-PR: area under the precision-recall curve
AUC-ROC: area under the receiver-operating-characteristic curve
BH: Benjamini-Hochberg
CAF: cancer-associated fibroblast
CF: cystic fibrosis
CNV: copy-number variation
CO: control
EMT: epithelial-mesenchymal transition
ER: estrogen receptor
FDR: false discovery rate
GMT: gene matrix transposed
GSEA: gene set enrichment analysis
HER2: human epidermal growth factor receptor 2
IFN: interferon
LLM: large language model
LOOCV: leave-one-out cross-validation
MNN: mutual nearest neighbours
MSigDB: Molecular Signatures Database
PC1: first principal component
PCA: principal component analysis
ROMA: Representation and Quantification Of Module Activity
scRNA-seq: single-cell RNA sequencing
SVD: singular value decomposition
TA: transit-amplifying
TNBC: triple-negative breast cancer
UMAP: uniform manifold approximation and projection.

## Declarations

### Ethics approval and consent to participate

Not applicable. This study analyzed only previously published, publicly available, de-identified human datasets; ethical approval and informed consent were obtained by the investigators of the original studies.

### Consent for publication

Not applicable.

### Availability of data and materials

All datasets analyzed in this study are publicly available: the PBMC IFN-*β* dataset (GEO GSE96583) [30], the cystic fibrosis airway atlas (GEO GSE150674) [31], the breast cancer atlas (GEO GSE176078) [32], the lung adenocarcinoma dataset of Maynard et al. [33] (NCBI Bio-Pro ject PRJNA591860), and the intestinal-organoid atlas of Capeling et al. [40] (GEO GSE313368).

The datasets generated during the current study are deposited in the Zenodo repository under two records. The ten pre-generated simulation replicates per regime, together with the corresponding gene-set definitions and all generative parameters, are available at https://doi.org/10.5281/zenodo.21699535 [49]. The derived intermediates the scANVI-integrated cystic fibrosis object underlying Fig. 4 and Supplementary Fig. S5, the inferCNV-annotated lung adenocarcinoma subset underlying Fig. 7 and Supplementary Figs. S8 and S9, the batch-aware scROMA outputs for the breast cancer atlas, and the tumor-only scROMA outputs for the lung dataset — are available at https://doi.org/10.5281/zenodo.21702006 [50]. All remaining derived intermediates, comprising the per-pathway scROMA outputs, per-cell pathway-activity score matrices, figure source-data tables and the complete per-replicate benchmark metrics, are versioned in the analysis repository below and archived with it.

### Software and analysis code

- **Project name:** scROMA
- **Project home page:** https://github.com/sysbio-curie/scroma [45]
- **Archived version:** version 0.3.2, archived at Zenodo [46]
- **Analysis code:** https://github.com/altyn-bulmers/scroma-reproducibility [47], archived at Zenodo [48]
- **Operating system(s):** Platform independent
- **Programming language:** Python
- **Other requirements:** Python ≥ 3.10; Scanpy, AnnData, scikit-learn, SciPy, NumPy, joblib
- **License:** BSD 3-Clause
- **Any restrictions to use by non-academics:** None

Analysis scripts reproducing every figure are provided in the reproducibility repository; the large intermediates held in the Zenodo data record [50].

### Competing interests

The authors declare no competing interests.

### Funding

A.Z. was supported by a doctoral contract from Ecole Doctorale 474 FIRE, Universite Paris Cite, Learning Planet Institute. This work was granted access to the HPC resources of IDRIS under the allocation AD011016305 made by GENCI (Grand Equipement National de Calcul Intensif).

### Author contributions

A.Z. L.M. conceived the study, E.B. contributed to the study design. A.Z. developed the scROMA methodology and software, performed the analyses, and drafted the manuscript. E.B. and L.M. supervised the project. M.N. and V.L. contributed to discussion and interpretation of the results and critically revised the manuscript. All authors read and approved the final manuscript.

## Acknowledgements

The authors thank the members of the Computational Systems Biology of Cancer laboratory (INSERM U1331), Institut Curie, for helpful discussions and feedback throughout this work. We are grateful to Anand D. Jeyasekharan and the members of his laboratory at the Cancer Science Institute of Singapore (CSI), National University of Singapore, for helpful discussions.

## Use of AI and AI-assisted technologies

The authors’ use of a large language model as an assistive tool for language editing and code refactoring is described in Methods, Section 5.7. No AI tool is listed as an author; the authors take full responsibility for the content of this manuscript.

## Notes

https://doi.org/10.5281/zenodo.21699535

https://doi.org/10.5281/zenodo.21702006

### Competing Interest Statement

The authors have declared no competing interest.

