## Supplementary material for "scROMA: batch-aware pathway-activity inference and a ground-truth simulation framework for single-cell transcriptomics": Figure S1

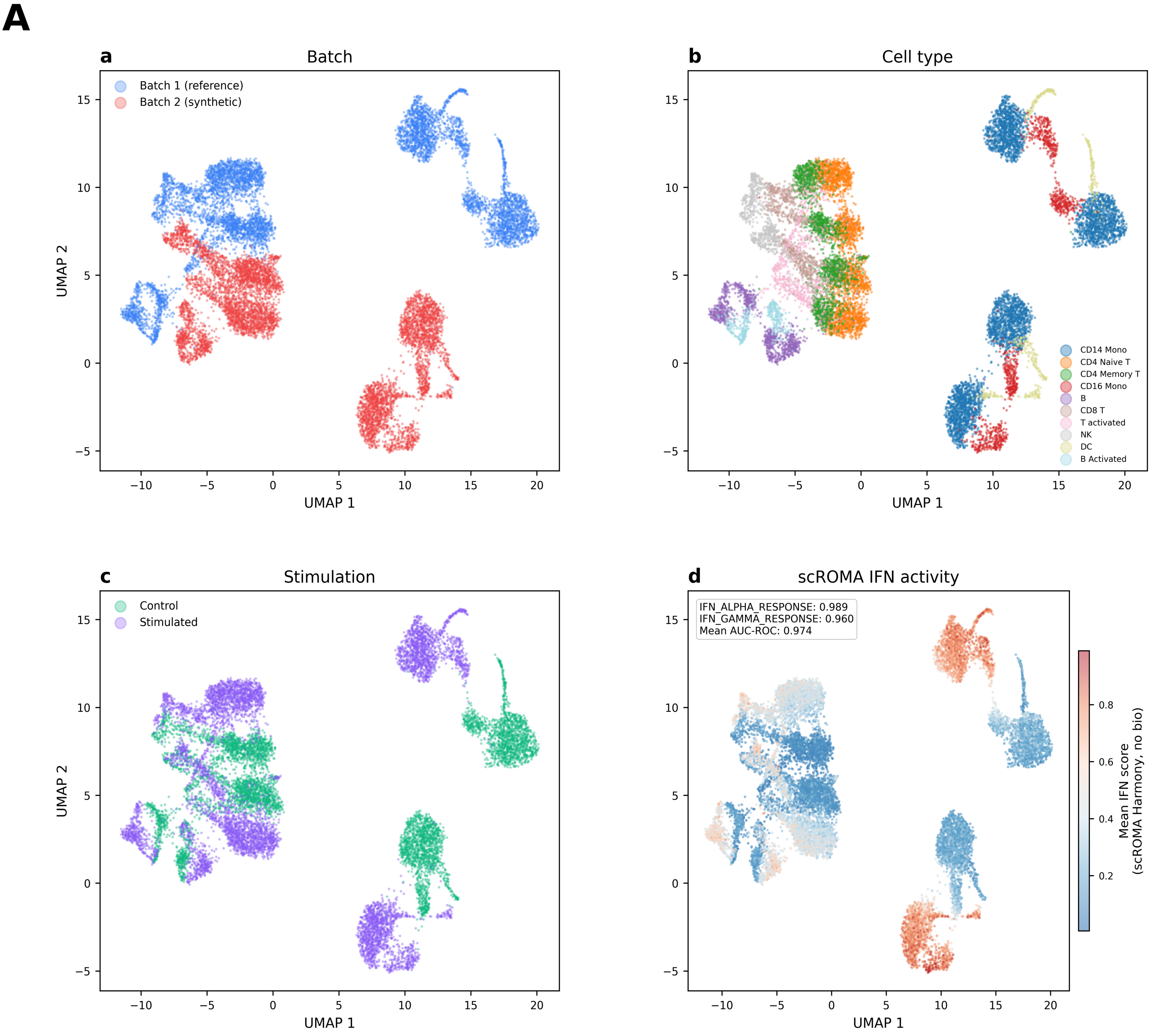


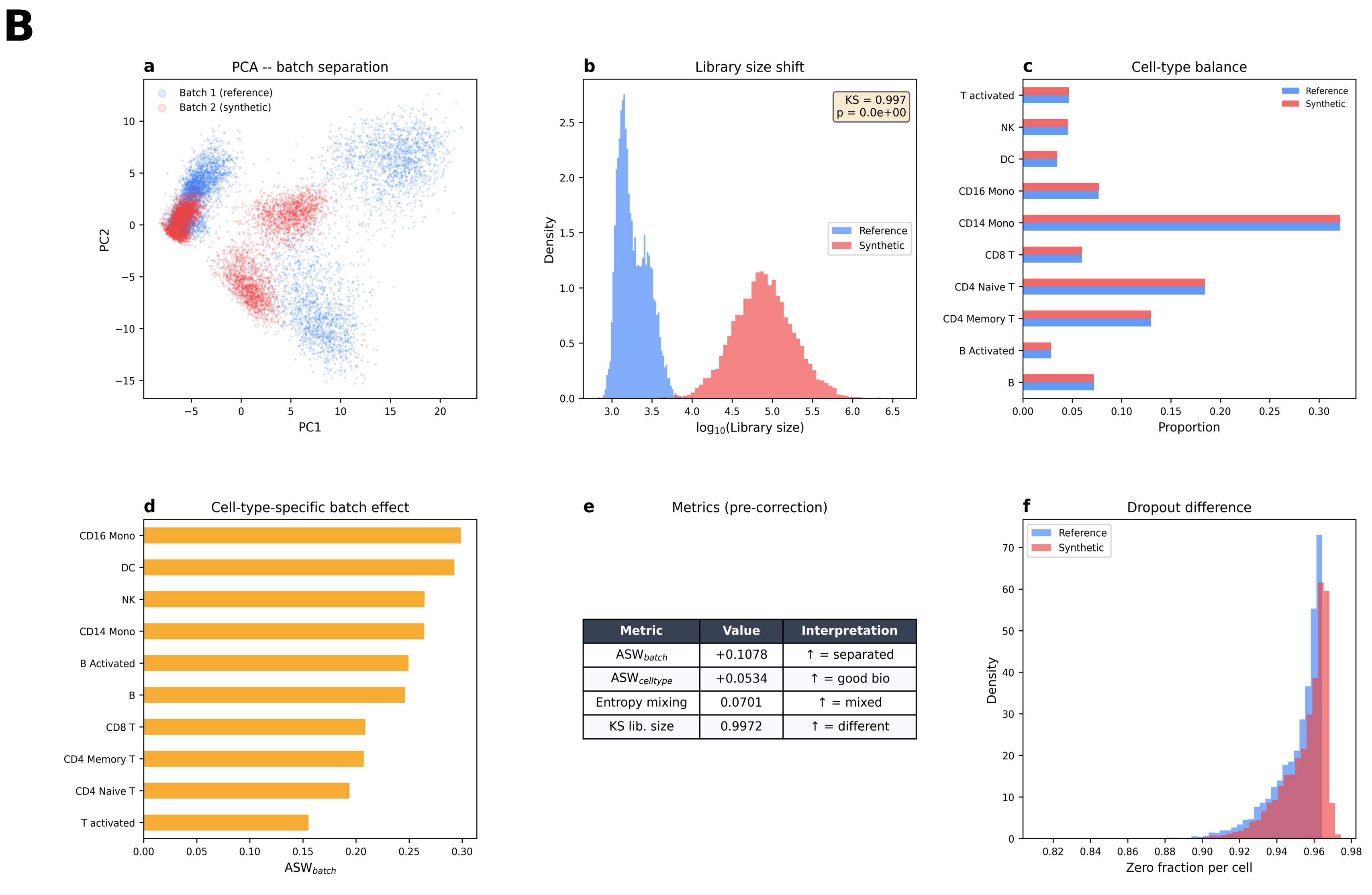


| Metric | Value | Interpretation |
|---|---|---|
| $ASW_{batch}$ | +0.1078 | ↑ = separated |
| $ASW_{celltype}$ | +0.0534 | ↑ = good bio |
| Entropy mixing | 0.0701 | ↑ = mixed |
| KS lib. size | 0.9972 | ↑ = different |
