## Supplementary figures and images for "scROMA: batch-aware pathway-activity inference and a ground-truth simulation framework for single-cell transcriptomics"

### Figure S2

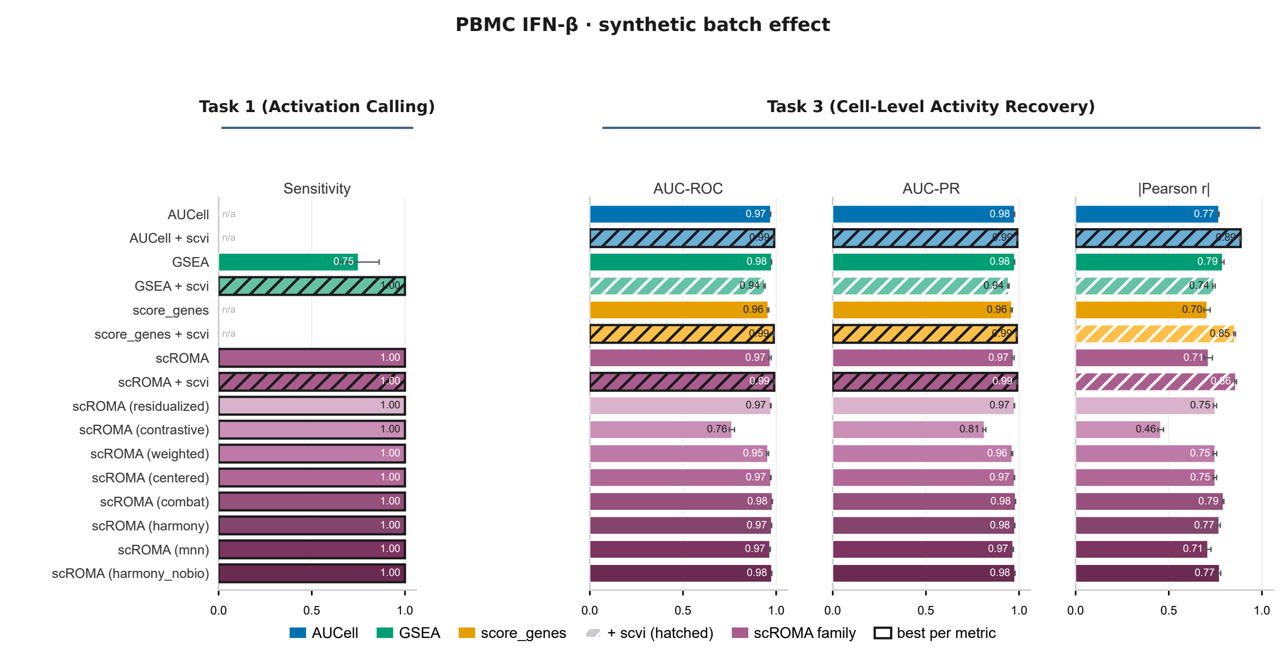

### Figure S3

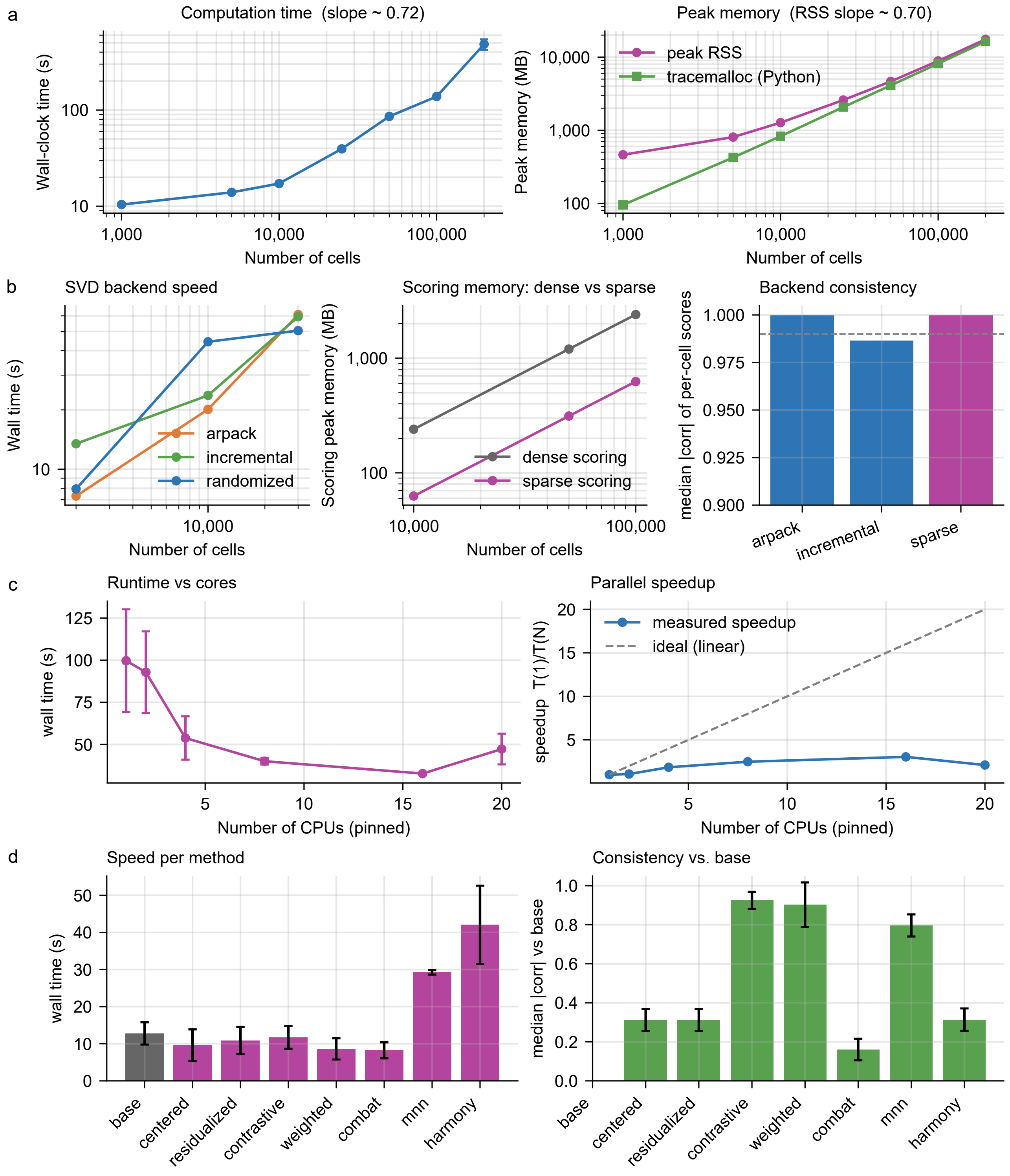

### Figure S4

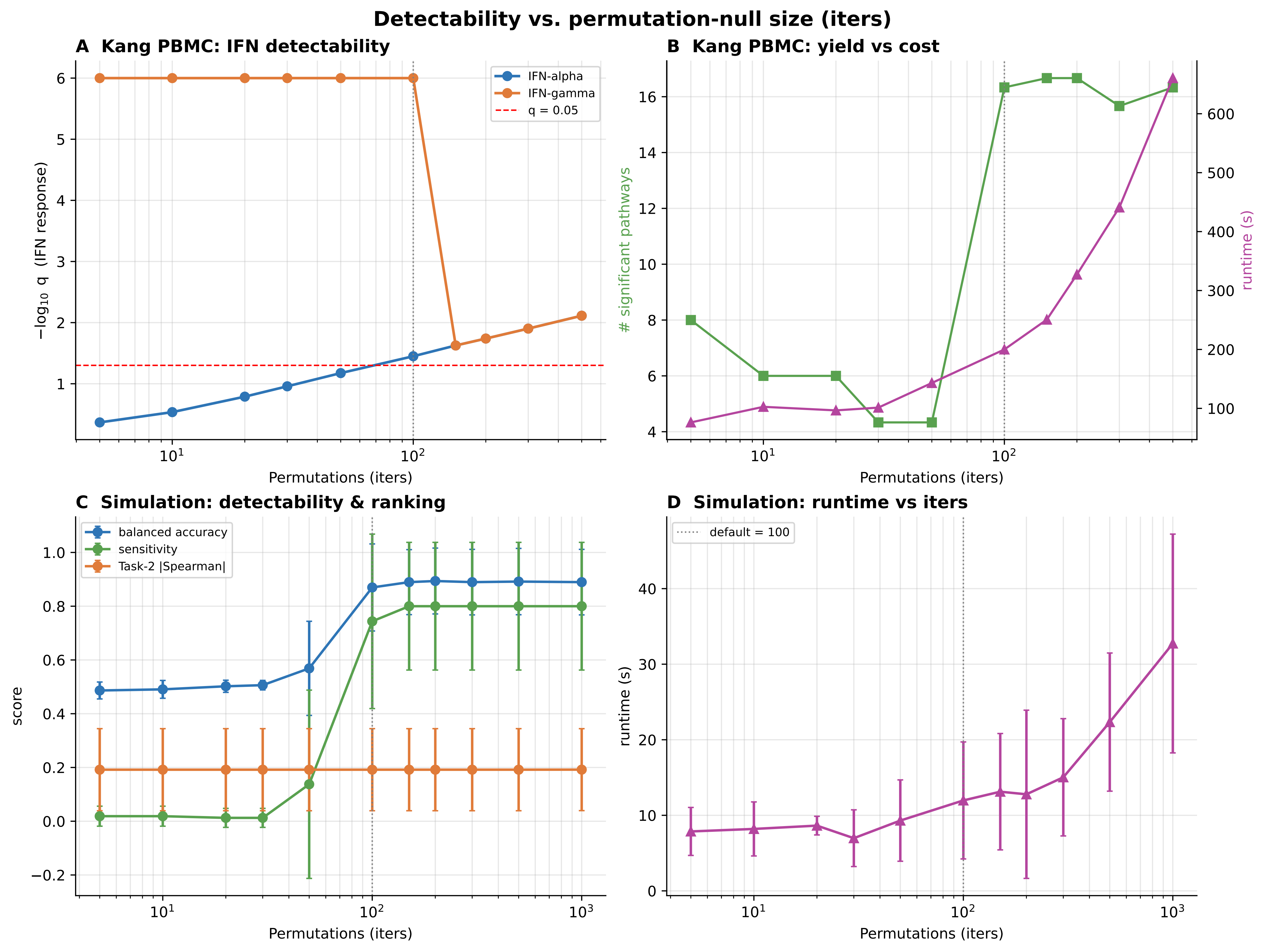

### Figure S5

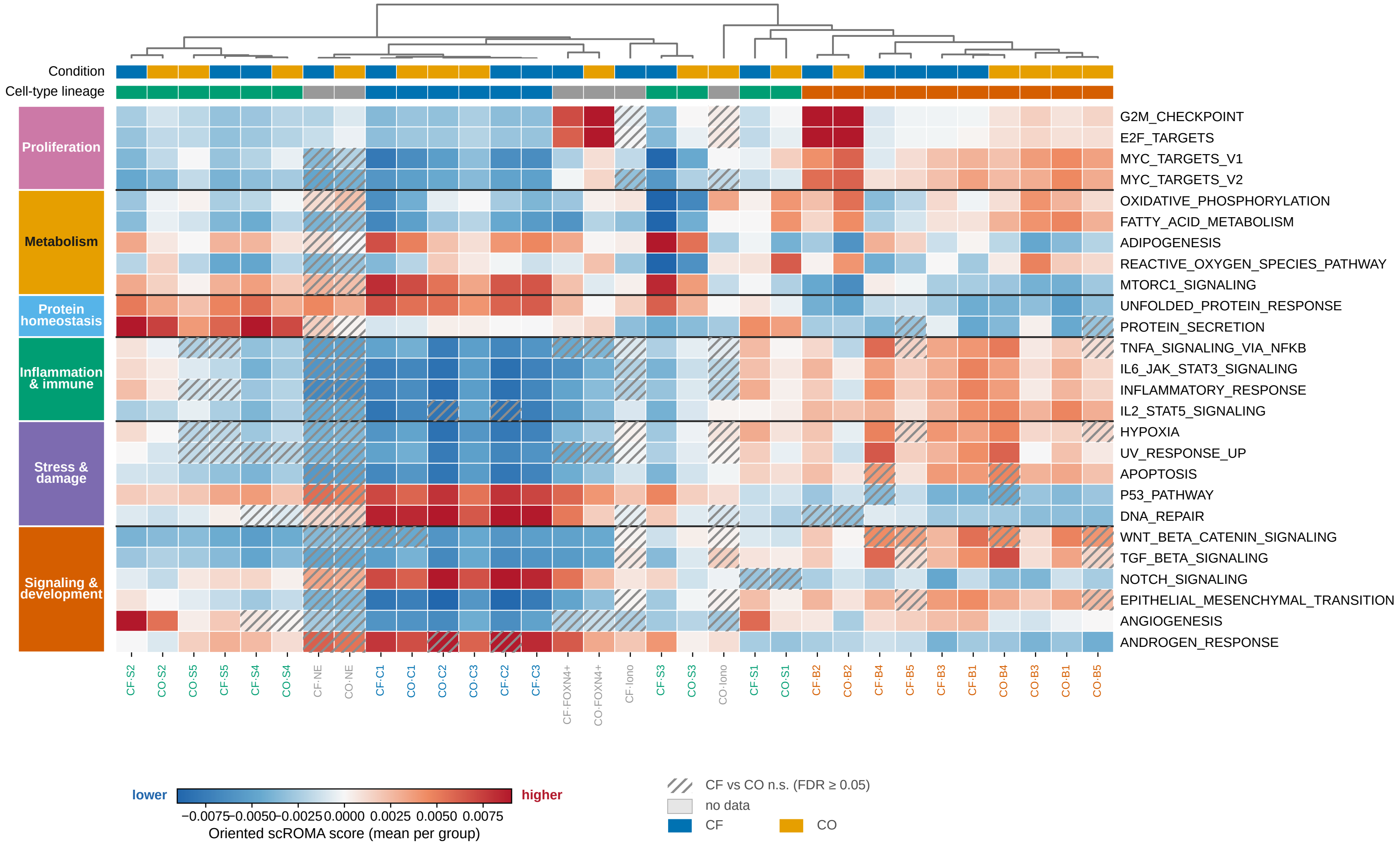

### Figure S6

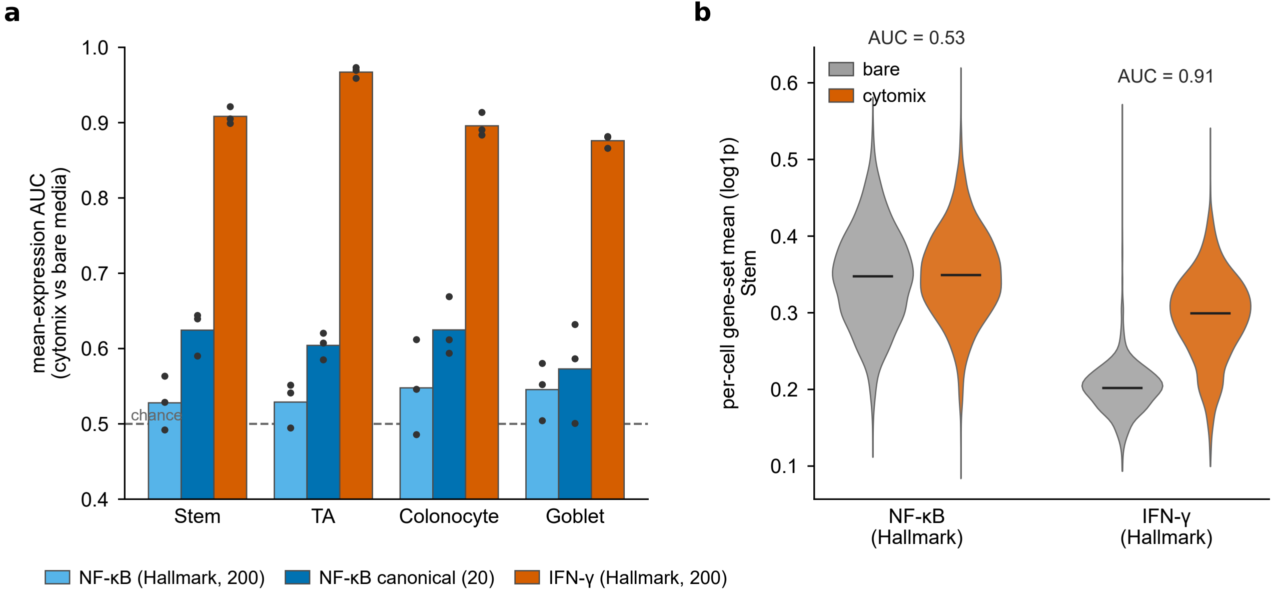

### Figure S7

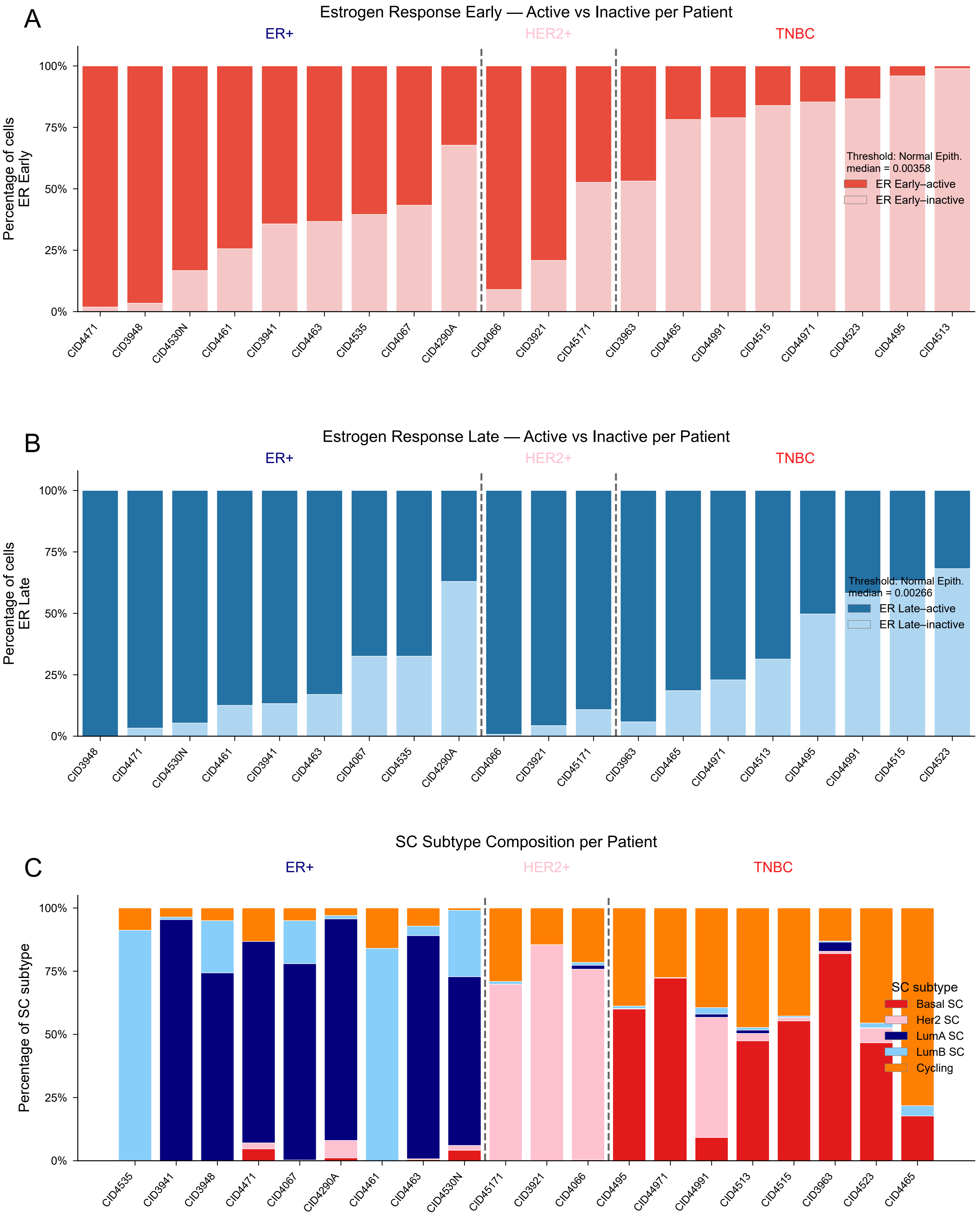

### Figure S8

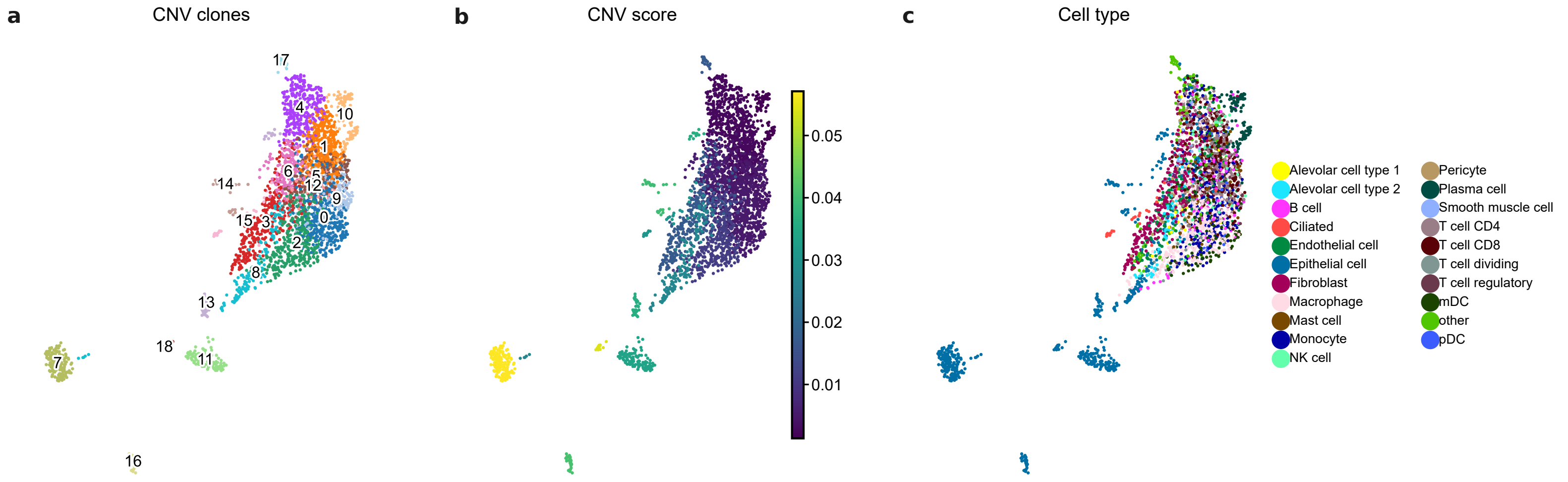

### Figure S9

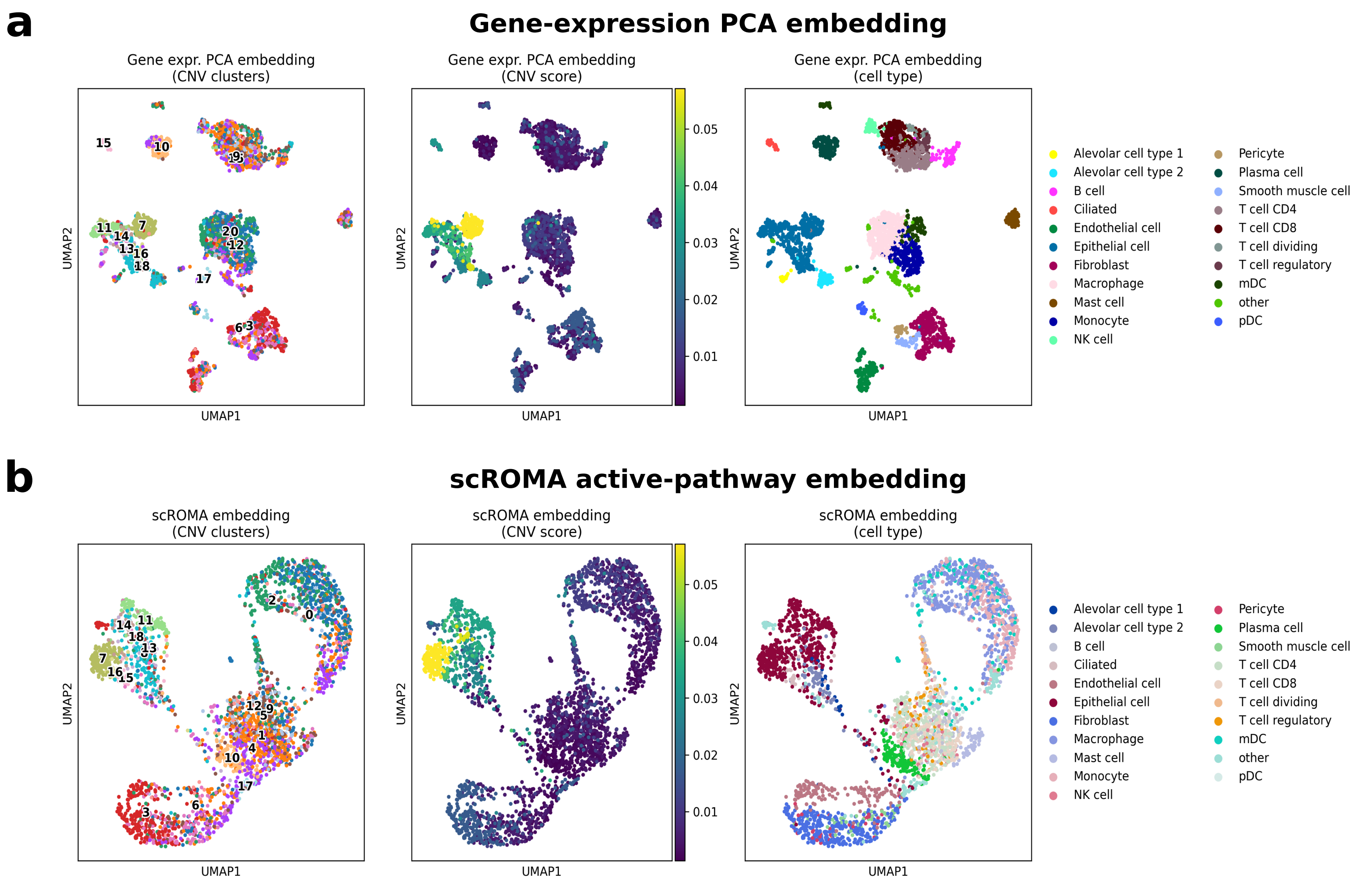

### Figure S10

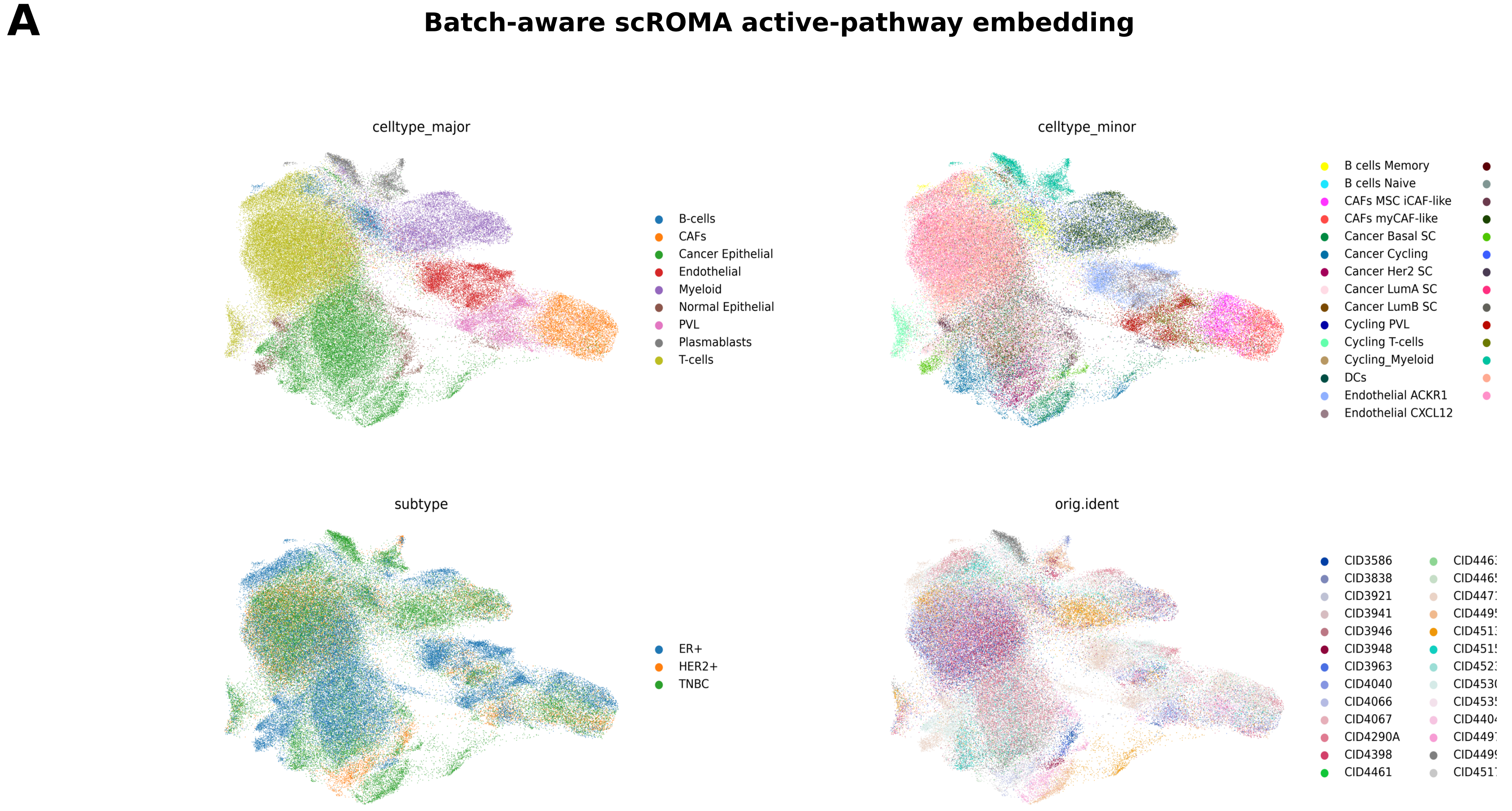

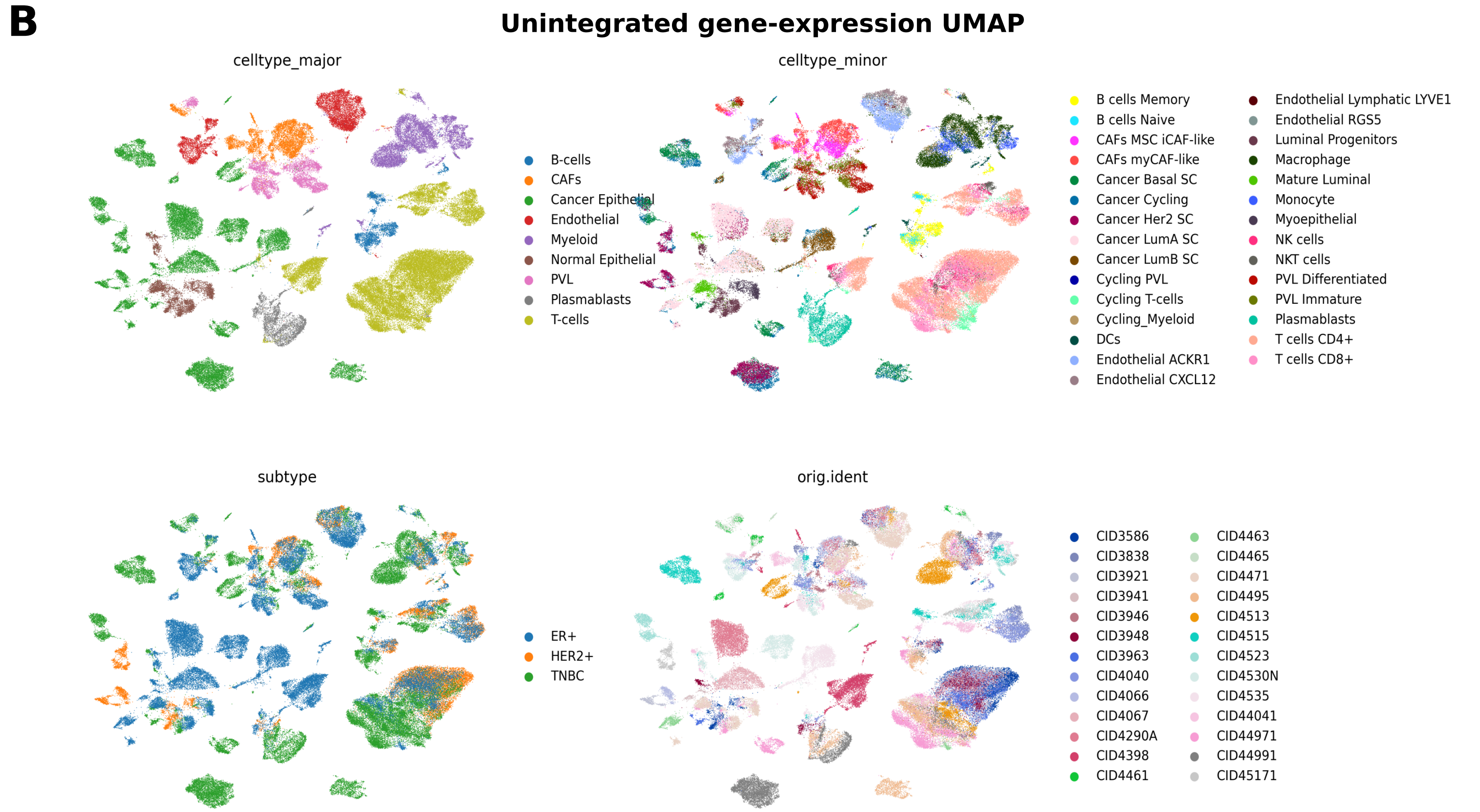
